# The chloroplast ribonucleoprotein CP33B quantitatively binds the *psbA* mRNA

**DOI:** 10.1101/2020.02.11.944249

**Authors:** Marlene Teubner, Benjamin Lenzen, Lucas Bernal Espenberger, Janina Fuss, Jörg Nickelsen, Kirsten Krause, Hannes Ruwe, Christian Schmitz-Linneweber

**Author notes:** Current address: Institute of Clinical Molecular Biology, Christian Albrechts University Kiel, Am Botanischen Garten 11, 24118 Kiel, Germany. corresponding author: Christian Schmitz-Linneweber.

## Abstract

Chloroplast RNAs are stabilized and processed by a multitude of nuclear-encoded RNA binding proteins, often in response to external stimuli like light and temperature. A particularly interesting RNA based regulation occurs with the *psbA* mRNA, which shows light-dependent translation. Recently, the chloroplast ribonucleoprotein CP33B was identified as a ligand of the *psbA* mRNA. We here characterized the interaction of CP33B with chloroplast RNAs in greater detail using a combination of RIP-chip, quantitative dot-blot, and RNA-Bind-n-Seq experiments. We demonstrate that CP33B prefers *psbA* over all other chloroplast RNAs and associates with vast majority of the *psbA* transcript pool. The RNA sequence target motif determined *in vitro* does not fully explain CP33B’s preference for *psbA*, suggesting that there are other determinants of specificity *in vivo*.

## Introduction

The maturation and translation of chloroplast RNAs depends on numerous RNA binding proteins (RBPs). With few exceptions, all RBPs involved in chloroplast RNA metabolism are encoded in the nucleus and are post-translationally imported into plastids. The largest family of RBPs in chloroplasts are the pentatricopeptide repeat proteins (PPR proteins), which interact specifically with one or few chloroplast transcripts (Barkan and Small, 2014). In addition to PPR proteins, several smaller RBP families exist in the chloroplast, including the family of chloroplast ribonucleoproteins (cpRNPs), whose members are extremely abundant, bind multiple mRNAs, and which are regulated in response to various biotic and abiotic signals (Ruwe et al., 2011).

cpRNPs are characterized by their conserved domain structure. An N-terminal chloroplast import signal (transit peptide), which is cleaved off after transport into the chloroplast, is followed by an acidic domain and two RNA recognition motif (RRM) domains. cpRNPs are able to interact with different nucleic acids (ssDNA, dsDNA and RNA; Li and Sugiura, 1990; Li and Sugiura, 1991), but the strongest association *in vitro* is with ribonucleic acids (Ye and Sugiura, 1992). *In vitro* and *in vivo* interactions with specific plastid mRNAs were demonstrated (Nakamura et al., 1999), cumulating in transcriptome-wide binding studies that showed a broad range of target mRNAs for various cpRNPs (Kupsch et al., 2012; Teubner et al., 2017; Watkins et al., 2019). rRNAs and intron-less tRNAs are not or only weakly bound (Nakamura et al., 1999; Nakamura et al., 2001; Kupsch et al., 2012). Since the cpRNPs do not co-fractionate with polysomal RNAs (Lisitsky et al., 1995; Nakamura et al., 1999; Teubner et al., 2017), they are mainly attributed a function prior to translation within posttranscriptional processes.

Prediction algorithms for subcellular localization and shotgun proteome analysis identified all ten Arabidopsis members in the chloroplast (summarized in Ruwe et al., 2011). Fluorescence microscopy of GFP fusion proteins confirmed the chloroplast localization (Raab et al., 2006; Fu et al., 2007; Xu et al., 2013; Teubner et al., 2017). Within the chloroplasts, the stroma is the main destination of cpRNPs, with small amounts also being associated with thylakoids. This was proven by immunological analyses for the five cpRNPs from tobacco (Nakamura et al., 1999).

The expression of cpRNPs is regulated by various external and internal signals. Especially light leads to an accumulation of cpRNPs (summarized in Ruwe et al., 2011). In general, cpRNPs are involved in a variety of posttranscriptional processes, including 3’-end processing of RNAs (Schuster and Gruissem, 1991), RNA editing (Hirose and Sugiura, 2001; Tillich et al., 2009), RNA splicing (Kupsch et al., 2012), and RNA stabilization (Nakamura et al., 2001; Kupsch et al., 2012; Teubner et al., 2017). Some of these processes are modulated by cpRNPs in response to environmental cues and several cpRNPs have been implicated in different acclimation and stress responses (Ruwe et al., 2011; Kupsch et al., 2012; Vargas-Suarez et al., 2013; Xu et al., 2013). Such a multi-level and far-reaching regulation by multiple external and internal stimuli is unknown for most other chloroplast RBPs, including PPR proteins. cpRNPs are thus considered as prime candidates for post-transcriptional regulators of plastid gene expression (Tillich et al., 2010).

A particular interesting case of chloroplast gene regulation is light-induced translation of *psbA*, which codes for the D1 protein, the core subunit of photosytem II (reviewed in Nickelsen et al., 2014; Sun and Zerges, 2015; Chotewutmontri and Barkan, 2018; Zoschke and Bock, 2018). D1 is constantly damaged, most pronouncedly by excess light and other unfavourable conditions, i.e. cold. As a consequence, D1 is constantly synthesized for repair of PSII (Sundby et al., 1993; Tyystjarvi and Aro, 1996; Nishiyama and Murata, 2014). Moreover, regulated D1 synthesis for de novo biogenesis of PSII during cell growth requires additional regulatory levels of *psbA* mRNA translation. Consistently, a number of proteins have been co-purified with the *psbA* mRNA in Chlamydomonas, spinach, Arabidopsis and maize, and or have been identified by genetic analyses (Danon and Mayfield, 1991; Alexander et al., 1998; Shen et al., 2001; Ossenbuhl et al., 2002; Schult et al., 2007; Link et al., 2012; Bohne et al., 2013; McDermott et al., 2019; Williams-Carrier et al., 2019). Among the proteins co-precipitating with the *psbA* mRNA was CP33B (AT2G35410; Watkins et al., 2019). In contrast to other cpRNPs, CP33B appears to have a clear preference for the *psbA* mRNA and does show comparatively little binding to other mRNAs (Watkins et al., 2019). We here analyzed the binding preference of CP33B in more detail and identified a main target sequence motif *in vitro*.

## Results

### CP33B localizes to the chloroplast

Arabidopsis CP33B is predicted to be a chloroplast protein based on algorithms such as Predotar and TargetP (Ruwe et al., 2011). This is supported by the recent finding that a maize orthologue of CP33B was isolated from chloroplast stroma by precipitating the *psbA* mRNA (Watkins et al., 2019). We analyzed the location of Arabidopsis CP33B experimentally by assaying cell fractions immunologically. Chloroplasts were isolated from 14 day old wild type plants and separated into membranes and soluble proteins (= stroma). The membranes were washed several times and a part was solubilized with the anionic detergent sodium deoxycholate (0.5%) (**Fig. 1**). From each fraction, equal volume portions were separated by SDS PAGE. A PsaD antibody, which detects a peripheral subunit of photosystem I, was used as membrane marker. No signal is detected for PsaD in the stroma fractions, a strong signal in the membranes and only a very weak signal in the detergent-treated membranes. Thus, not even peripheral membrane proteins such as PsaD are released from the membrane by the treatment. The detection of RbcL, the large subunit of the ribulose 1,5 bisphosphate carboxylase/oxygenase (RuBisCO) located in the stroma, serves as a marker for the stroma fraction. The Western analyses showed that the majority CP33B, like CP33A is found in the stroma. In contrast to CP33A that showed only a very week membrane association, CP33B was also present to a considerable degree in the membrane fraction is found in the stroma. (Fig. 1). We quantified the CP33B signals and found that about 75% of CP33B is found in the stroma and about a quarter in the membrane fraction. Since CP33B can be easily dissolved from the membrane fractions with the mild detergent sodium deoxycholate, the membrane-bound part of CP33B is not integrated into the membranes, but is likely to be bound peripherally, via weak interactions to the membranes. Overall, the largest proportion of CP33B localizes in the stroma, in line with all previous analysis of cpRNP proteins.

**Figure 1:**
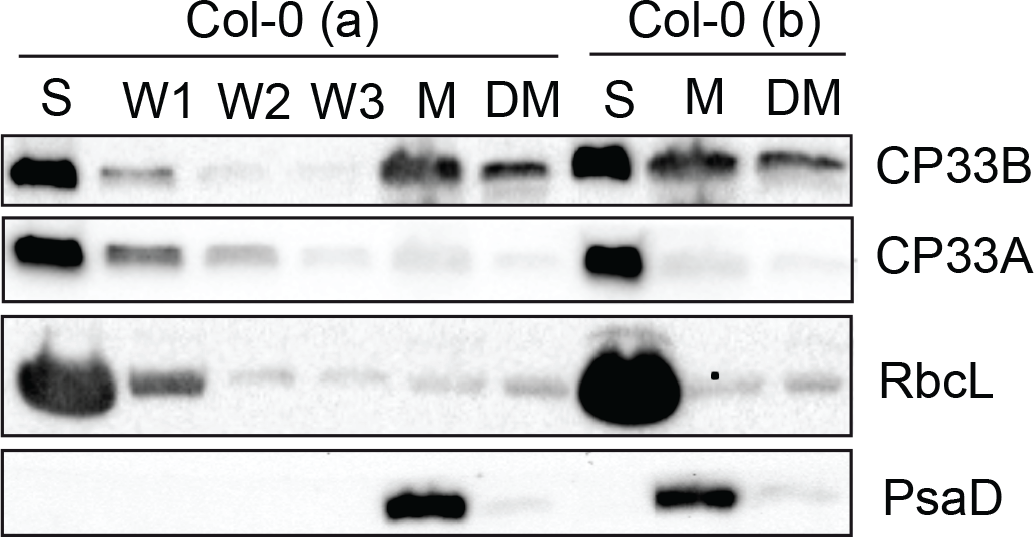
CP33B localizes to stroma and membrane fraction of chloroplasts Chloroplasts of two wild type replicates (Col-0 a and b) were separated into stroma (S) and membranes (M). Membranes were washed 5 times. A membrane aliquot was treated with 0.5% sodium deoxycholate and the supernatant was harvested after treatment (DM). From each fraction, including the first three washing steps (W1, W2, W3) of Col-0 (a), equal volume fractions were separated electrophoretically by SDS PAGE and transferred to a nitrocellulose membrane. Besides the immunological detection of CP33A (antibody peptide a) and CP33B, antibodies for RbcL (large subunit of RuBisCO (ribulose 1.5 bisphosphate carboxylase/oxygenase)) and PsaD (subunit of photosystem I) served as stroma and membrane markers, respectively. The controls (RbcL, CP33A, PsaD) are reprinted with permission from Wiley; original figure 2c in Teubner et al. (2017).

**Figure 2:**
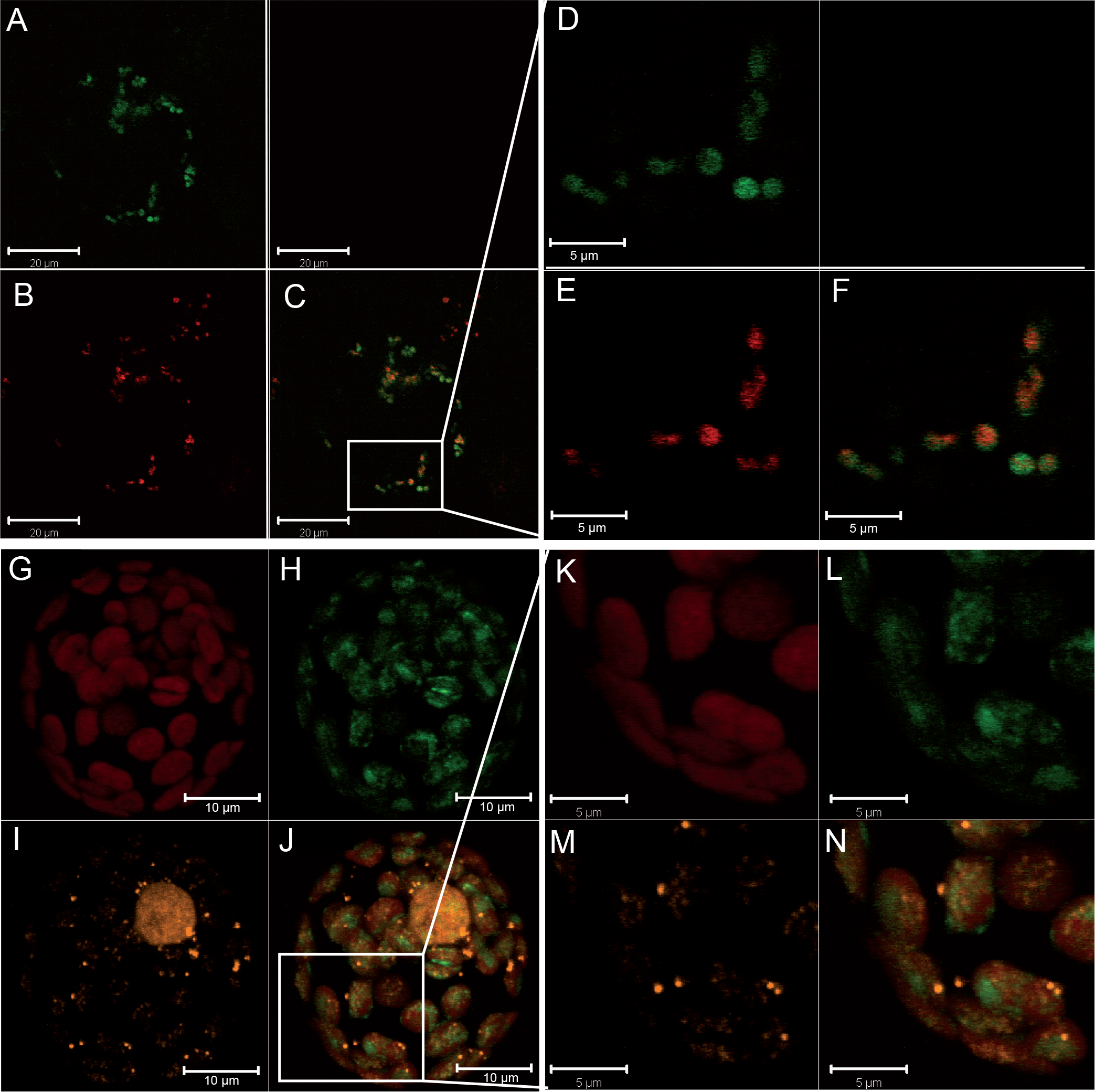
CP33B is not associated with nucleoids. A-F) Images of CP33B:GFP transformed Arabidopsis protoplasts. D, E, and F are enlargements of the white box area of A, B, and C respectively. GFP fluo-rescence (A and D, green); chlorophyll autofluorescence (B and E, red); super-imposition of images A and B (in C) and D and E (in F), respectively. G-N) Images of co-transfection experiments of CP33B:GFP and PEND:dsRed. K-N are enlargements of the white box area of G-J. Chlorophyll autofluorescence (G and K, red); GFP fluorescence (H and L, green); dsRed fluorescence (I and M, orange); superimposition of images G-I (in J) and K-M (in N), respectively. The white bars represent 20 μm, 10 μm and 5 μm, as indicated.

The stroma localization of CP33B was further analyzed by fluorescence microscopy. For this, the coding sequence of CP33B was fused with a GFP tag and transiently expressed in protoplasts. The GFP fluorescence signal is mainly diffusely distributed in chloroplasts, with a partial overlap with chlorophyll autofluorescence (**Fig. 2 A-F**). RBPs have been found to co-fractionate with nucleoid preparations (Majeran et al., 2011)and thus, we decided to test for nucleoid association as well. To analyze a potential co-localization of CP33B with nucleoids, a co-transfection was performed with a vector expressing PEND:dsRed. PEND binds to plastid DNA and can therefore be used as a marker for nucleoids (Terasawa and Sato, 2005). The fluorescence images show little or no overlap of CP33B GFP and the PEND dsRed signal (**Fig. 2 G-N**), which makes an association of CP33B with nucleoids unlikely. In sum, CP33B localizes predominantly to the chloroplast stroma with a minor fraction of CP33B attached to membranes.

### CP33B has a preference for the *psbA* mRNA over other chloroplast transcripts

Previously, we had shown by RNA-co-immunoprecipitation and next generation sequencing (RIP-Seq) that CP33B associates with a number of chloroplast mRNAs, but has a strong preference for *psbA* (Watkins et al., 2019). We validated this finding using an alternative detection technique, that is, microarray hybridization (RIP-chip; Schmitz-Linneweber et al., 2005). RIP-chip has the advantage over RIP-Seq that the co-precipiated RNA is directly labelled using a chemically activated dye without any further enzymatic steps. This avoids any experimental bias potentially introduced via enzymatic steps like reverse transcription, linker ligation, and PCR, or RNA size selections commonly used during library preparation for deep sequencing. Using the same antibody raised against a CP33B-specific peptide as in the RIP-Seq approach, we precipitated CP33B from chloroplast stroma. In line with our previous efforts, we found a clear preference of CP33B for the *psbA* mRNA (**Fig. 3A**). The microarray used for these RIP-chip analyses consisted of PCR-generated probes that are mostly more than 500 nt in length. We next performed a fine-mapping analysis using a previously described oligonucleotide-based microarray (Zoschke et al., 2010) to confirm our findings. The oligo-RIP-chip verified the exceptional strong enrichment of *psbA* mRNA in CP33B precipitations (**Fig. 3B**). Enrichment was also seen for other mRNAs, but to a lesser degree (**Fig. S1**). In sum, there is exceptionally strong association of CP33B with the *psbA* mRNA.

**Figure 3:**
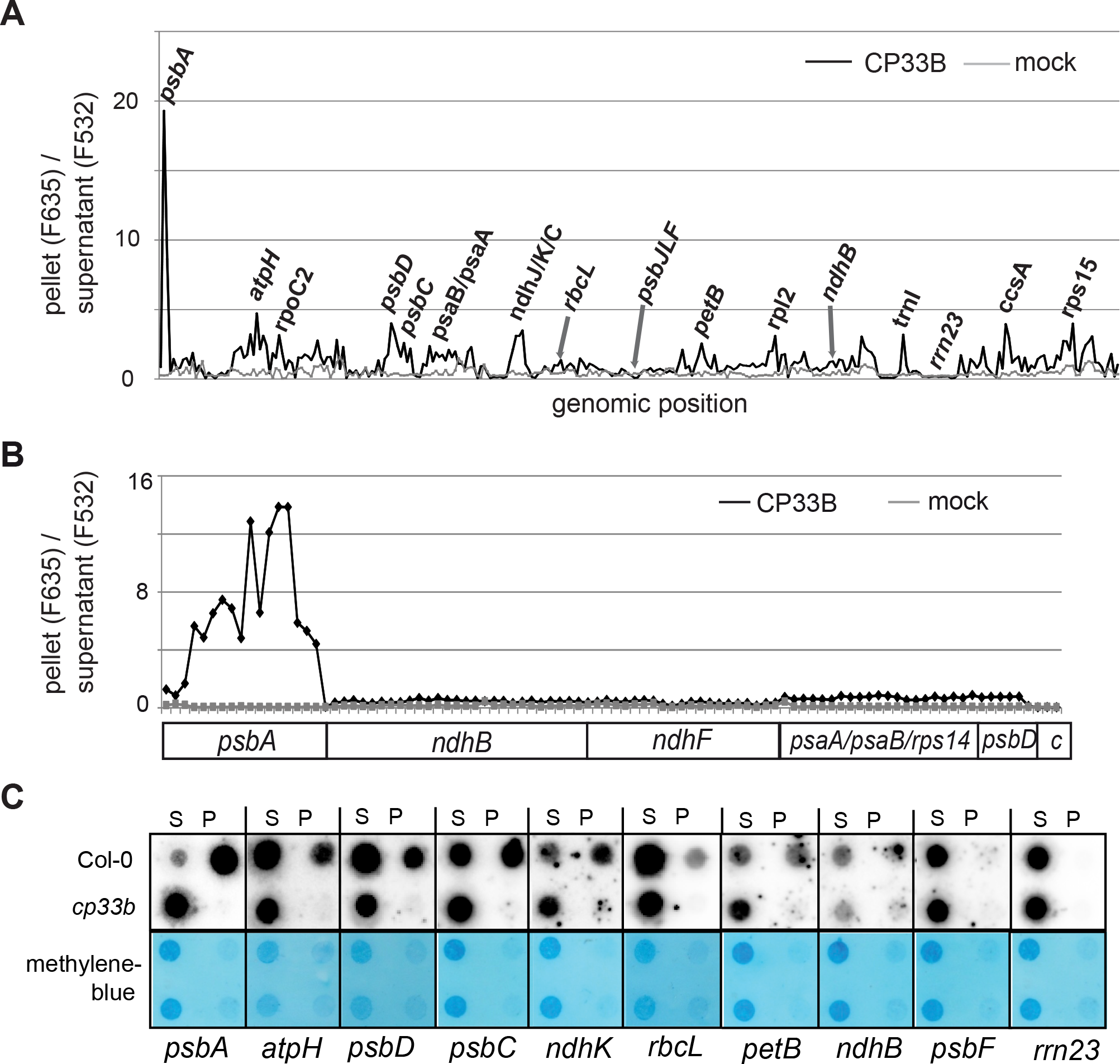
CP33B associated with multiple plastid transcripts, but foremost with the *psbA* mRNA A) RIP-chip analysis of RNA co-precipitated with CP33B using a microarray consisting of PCR products representing the entire Arabidopsis chloroplast genome with on average 1 kb fragments. Ratios of fluorescence signals from coprecipitated RNA (F635) and unbound RNA (F532) were plotted against the position on the chloroplast genome. Three biological replicates from CP33B immunoprecipitations and as control, two replicates of precipitations with the corresponding pre-immune serum and an IP on stroma of *cp33b* null mutants were performed. The precipitate/supernatant ratios (median (median of ratios)) were normalized to the sum of the median values of the ribosomal RNAs of the supernatants (median F532; Suppl. Table 1). Only selected peaks are labelled. B) Oligonucleotide RIP-chip analysis for the interaction of CP33B with a selection of chloroplast transcripts. CP33B immunoprecipitations were performed from wild-type stroma with anti CP33B (four biological replicates) and as a control with the corresponding pre-immune serum (three biological replicates). Shown are the median of ratios of fluorescence signals from coprecipitated RNA (F635) and unbound RNA (F532; sequences of oligonucleotides and data in Suppl. Tab. 2). Each plastid transcript, except for negative controls (c), is represented by several oligonucleotides (50 nt long; for a zoom-in see Suppl. Fig. 1). C) The CP33B immunoprecipitations were performed on wild type stroma and *cp33b* null mutants. Equal volumes of isolated RNA from precipitate (P) and supernatant fraction (S) were transferred onto nylon membranes and hybridized with different radiolabeled RNA probes. The methylene blue staining of the nylon membranes reflects the total RNA content of each fraction. The analysis was repeated twice for *psbA* and once more for *rbcL* and *rrn23* with similar results.

### CP33B quantitatively co-immunoprecipitates *psbA* mRNAs

The strong signal for *psbA* in CP33B RIP-chip experiments is remarkable given that this mRNA is the most abundant mRNA in chloroplasts. RIP-chip can however only give relative insights into RNA quantification, which prompted us to use dot-blot, a quantitative method to determine how much *psbA* mRNA is pulled down with CP33B. RNA from pellets and supernatants of CP33B precipitates from wt and from *cp33b* null mutant stroma preparations were blotted onto nylon membranes and probed with radiolabeled RNA probes. Importantly, equal fractions of supernatant and pellet volumes were blotted, which allows direct assessment of the efficiency of co-precipitation. Next to probes for several transcripts identified as targets of CP33B in RIP-chip analyses, we also used probes for two transcripts not enriched in RIP-chip assay, *psbF* and the 23S rRNA. The hybridization confirmed that these two control RNAs do not co-precipitate with CP33B (**Fig. 3C**). For all other transcripts tested, an accumulation in the precipitates from wild type stroma was detected, but not in the precipitates of the mutants. Importantly, the signal for *psbA* in the pellet was far stronger than the residual signal in the supernatant. The ratio of the pellet to supernatant signals was far higher for *psbA* than for any other coding region analyzed (74-fold enrichment for the top *psbA* probe versus 12-fold enrichment for the top *psbD* probe). Thus, the vast majority of *psbA* transcripts in the stroma was associated with CP33B. About half of the *psaC* and *ndhK* transcripts co-precipitated with CP33B. The *rbcL* mRNA showed the lowest enrichment of all transcripts analyzed. Overall, the dot blot analyses confirmed the results of the RIP chip experiments and demonstrated that a majority of all *psbA* transcripts is associated with CP33B.

### CP33B prefers an RNA sequence motif *in vitro*

We next asked, how the strong specificity of CP33B for *psbA* is determined. One explanation would be a target sequence motif that distinguishes the *psbA* mRNA from other chloroplast mRNAs. To test the RNA sequence preference of CP33B, we used the RNA-Bind-n-Seq (RBNS) assay (**Fig. 4A**). RBNS is an *in vitro* method that allows comprehensive mapping of RNA binding specificity (Lambert et al., 2014; Dominguez et al., 2018). It has considerable advantages over older *in vitro* methods like SELEX, which identify consensus motifs but is biased towards the highest affinity motifs (Campbell et al., 2012). By contrast, RBNS tests affinities to the full spectrum of possible RNA sequences in a high-throughput manner. Being an *in vitro* technique, RBNS tests the direct interaction of the protein with RNA targets and avoids common biases of RIP-Seq and RIP-chip, which can potentially also enrich for RNAs indirectly tethered to CP33B via protein partners.

**Figure 4:**
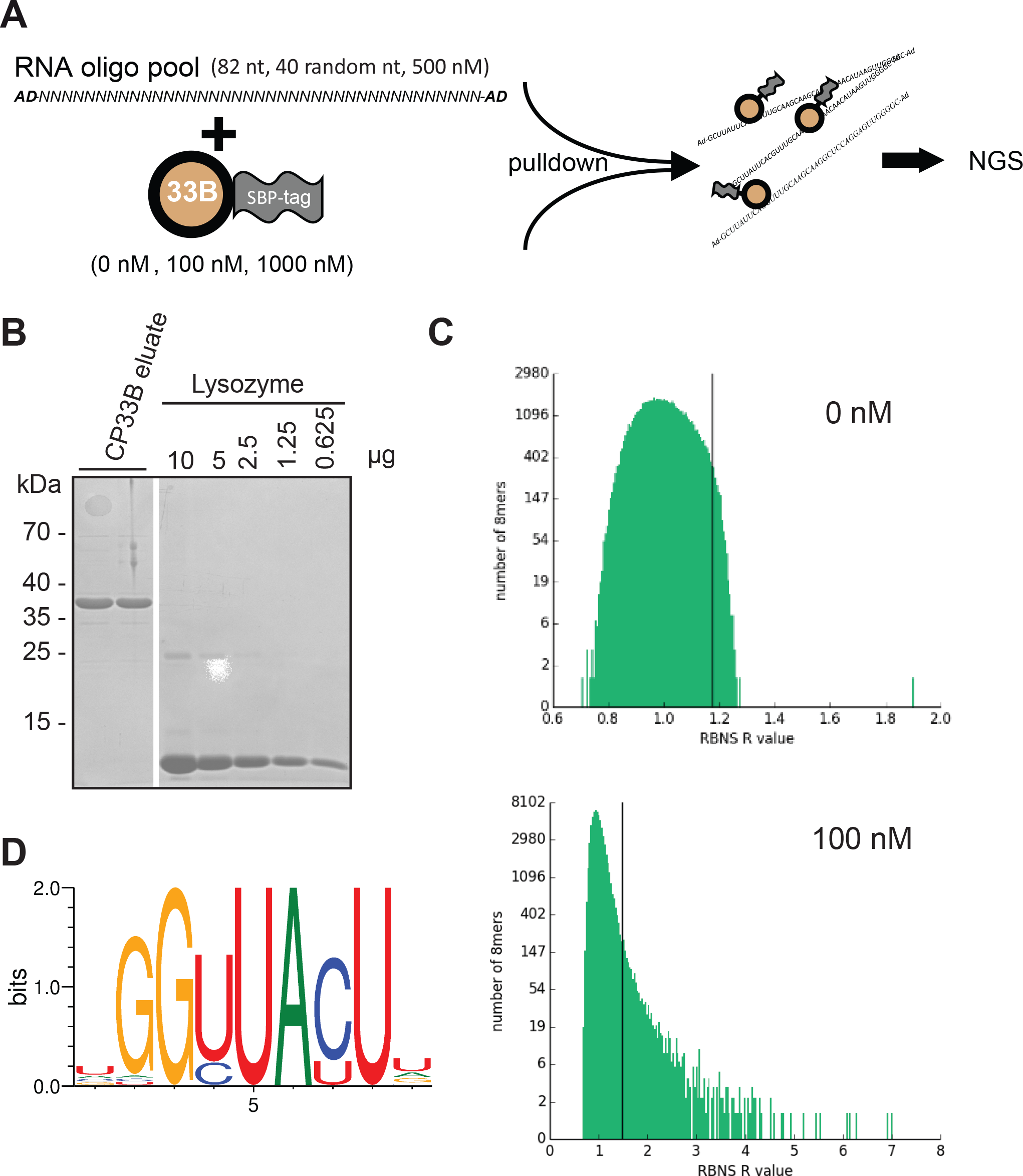
RBNS of CP33B A) Schematic overview of the RBNS experiment. B) Coomassie stain of an analytical PAGE to determine the amount of SBP:CP33B in two replicate eluates after protease-based removal of the GST tag. The white line indicates where lanes irrelevant for this analysis were removed. C) Stacked histogram showing the distribution of RBNS R values of all RNA 8mers in the CP33B experiment at a protein concentration of 0 and 100 nM, respectively. 8mers with R values that have z-scores equal or greater than two times the average are found on the right of the black lines. A log scale is used for the y axis. Please note the different scales for the X-axes between the two graphs. D) CP33B consensus motif generated from all 8mers with a z-score ≥ 3 in the 100 nM library (detailed rules on motif generation found in Dominguez et al. 2017).

As a prerequisite for this analysis, we expressed recombinant CP33B with a combined N-terminal GST:SBP tag, but without the known signal peptide for chloroplast targeting, and purified it using a Glutathione matrix. The GST tag is subsequently cleaved off using the PreScission Protease and remains on the purification column, while pure SBP-CP33B is eluted and quantified using a lysozyme standard (**Fig. 4B**). Two concentrations of recombinant SBP-CP33B (100 nM and 1000 nM) and a negative control (zero protein) were incubated with an input pool of random RNA 40mers. SBP-CP33B is pulled down using streptavidin-coated beads and the pulled down RNA is used for library amplification and subsequently sequenced. The input random RNA pool is also sequenced to account for potential compositional biases. For each of three protein concentrations (0, 100, and 1000 nM), enrichment (“R”) values were calculated for all *k*mers of selecetd lengths within the sequenced 40mers as the ratio of the frequency of the *k*mer in the sample pool to the frequency in the input pool. For CP33B, a large number of 6mer, 7mers, 8mers, and 9mer motifs had significant R values (as an example, the 8mer analysis in the 100 nM library is shown in **Fig. 4C**). In our experiments, the highest R values were observed at a 100 nM CP33B concentration. At a higher protein concentration, the ranking of top-enriched *k*mers remains similar, even though the overall observed R values are lower. This is an expected behavior in RBNS experiments since the top *k*mer targets are saturated at a certain protein concentration and secondary RNA targets become more co-precipitated as well, which lowers the overall R value. This observation and the ranking consistency between different libraries supports the validity of our application of RBNS to CP33B. For a more detailed analysis, we chose 8mers, since single RRM domains typically associate with 2-4 nucleotides and thus, the two RRMs of CP33B may associate with up to 8 nucleotides. All significant 8mers (with a Z-score ≥ 3) in the library with the highest enrichment (100 nM sample) were used to generates a consensus motif (details on this process in Dominguez et al., 2018), which reflects the binding preference of CP33B (**Fig. 4D**). We next asked, whether the RBNS results are reflecting the preference of CP33B for the *psbA* mRNA. However, when perfoming sequence searches for derivatives of the top motif, there were dozens of hits throughout the chloroplast genome, but only one hit in sense direction of the *psbA* coding region (**Suppl. Tab. 3**). Therefore, there must be other factors than simple sequence preferences *in vivo* that lead to the observed preference of CP33B for the *psbA* mRNA. Also, the conditions used in our RBNS analysis might not adequately reflect the *in vivo* situation, preventing us from the finding the true *in vivo* target sequence.

### Membrane-bound CP33B is also associated with chloroplast mRNAs

The RIP chip experiments shown in Fig. 3 were performed with stroma material as input. Given that a part of the CP33B pool is associated with membranes, we performed RIP-chip on isolated membrane fractions that were solubilized prior to immunoprecipitation. Overall, RNA enrichment in the membrane RIP chip is lower than in the stroma RIP-chip, which is indicated by the relatively small difference in enrichment values of CP33B-immunoprecipitation from wild-type membranes versus from control (*cp33b* mutant) membranes (**Fig. 5**). Despite the lower signal, the analysis reveals that CP33B associates with a multiple mRNAs also at membranes. Again, the top enriched mRNA is *psbA*, but the signal appears less dominant over secondary targets like *psbD/C, psaB/psaA, ndhJ/K/C, petB* and *atpF/H*. The 20 strongest enriched transcripts of the stroma RIP chip are also found among the strongest enriched mRNAs in the membrane RIP-chip (**Suppl. Tab. 1 / 4**). CP33B thus interacts with multiple and largely identical chloroplast mRNAs in the stroma as well as at membranes.

**Figure 5:**
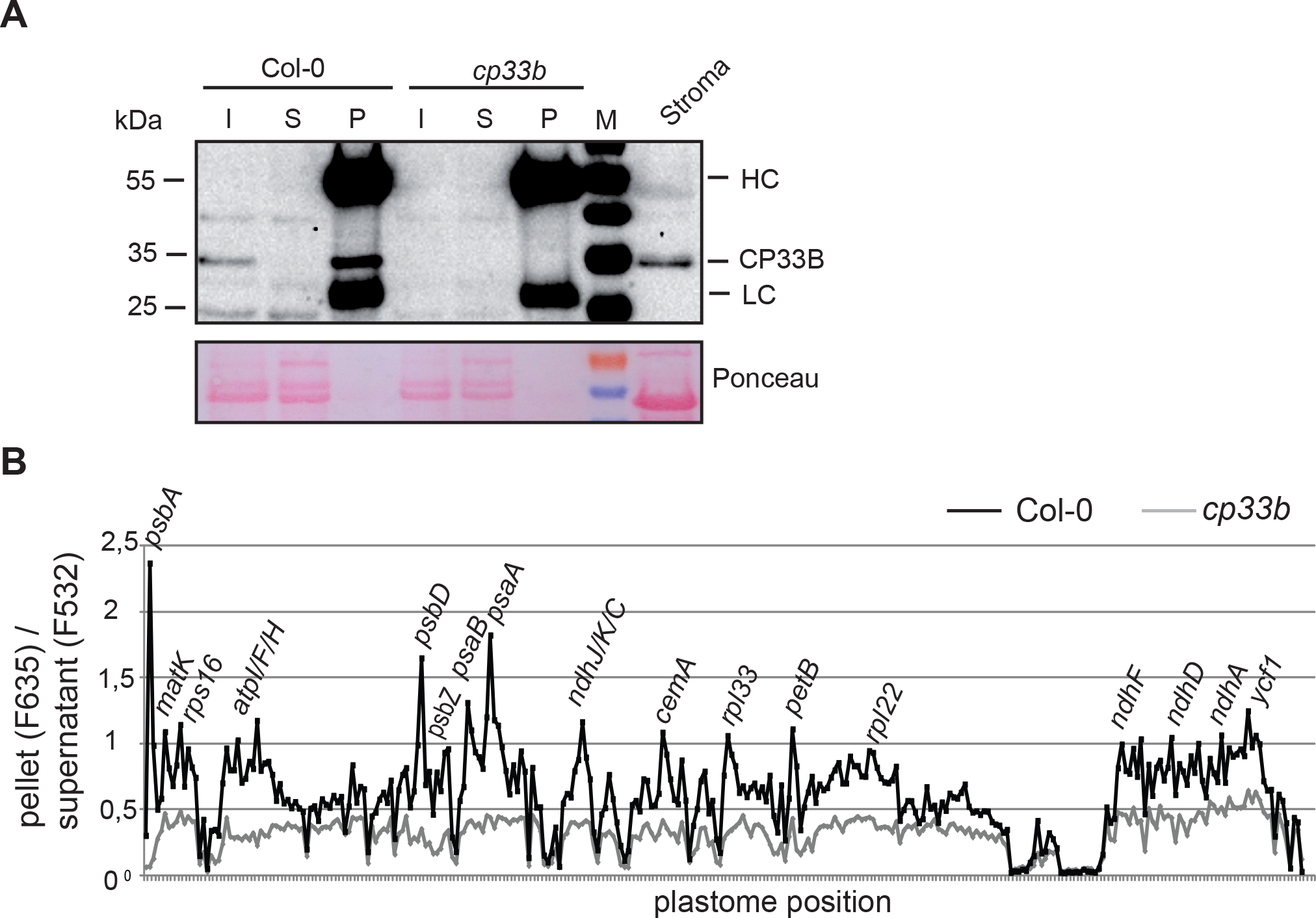
RNAs bound by membrane-localized CP33B A) Immunoprecipitation of CP33B from membranes Immunoblot analysis of protein fractions after immunoprecipiation of CP33B from solubilized membranes of wt and *cp33b* null mutants. Relative to volume of the input membrane fractions used (I) for the IPs, 1/20 of the supernatant (S) and 1/10 of the pellet (P) fractions were loaded. An aliquot of wt stroma from the same chloroplast preparation was analyed as well, which shows the depletion of RuBisCO in the membrane fractions compared to the stroma fraction by ponceau staining. Bands marked with HC and LC represent the heavy and light chains, respectively, of antibodies used for immunoprecipitation. B) RIP-Chip analysis of membrane-bound CP33B. RNA from pellet and supernatant fractions of CP33B immunoprecipitations and control reactions were analyzed by microarray hybridization analogously to stroma RIP-Chips shown in Fig. 3. The pellet/supernatant ratios (median (median of ratios)) were normalized to the sum of the median values of the ribosomal RNAs of the supernatants (median F532; Suppl. Tab. 4). Ratios of fluorescence signals from coprecipitated RNA (F635) and unbound RNA (F532) for IPs from wt and *cp33b* mutant membranes were plotted against the probe positions on the chloroplast genome.

## Discussion

### CP33B is a global RBP with a clear preference for and ability to sequester the *psbA* mRNA pool

The RNA binding spectrum of different RBPs can vary considerably. In higher plants, the PPR proteins, which are the most abundant family of RBPs in chloroplasts, have a small target spectrum of only one to a few transcript(s) (Barkan and Small, 2014). In contrast, the cpRNPs interact similar to their nucleo-cytoplasmic relative hnRNP A1 (Jean-Philippe et al., 2013) with a variety of RNAs, as shown for CP31A, CP29A, CP33A, and CP33B from Arabidopsis (Kupsch et al., 2012; Teubner et al., 2017).

In this study, the RNA targets of CP33B from *Arabidopsis thaliana* were investigated in greater detail. We confirmed that CP33B associates with a larger number of chloroplast mRNAs. Similar to CP31A and CP29A (Kupsch et al., 2012), CP33B showed little or no enrichment of ribosomal RNAs and tRNAs. Also, several of its minor targets, like *atpH*, *psbC/D*, and *psaA/B* are also targets of the three other cpRNPs analysed before (Kupsch et al., 2012; Teubner et al., 2017). Possibly, the four cpRNPs are all part of common ribonucleoprotein particles. This extended redundant target range may in part explain the lack of a detrimental phenotype of *cp29a*, *cp31a*, and *cp33b* mutants, respectively, under normal conditions, where loss of just one cpRNPs might be compensated by other members of the family.

Next to these commonalities in RNA targets, there are also striking differences for a selected low number of targets. Most strikingly, CP33B has a clear preference for a single mRNA, *psbA*, with an enrichment of 90% in precipitates. The *psbA* mRNA is no or only a weak target of CP33A, CP29A, and CP31A (Kupsch et al., 2012; Teubner et al., 2017). Vice versa, main targets of CP33A, like *psbF* or *rbcL*, of which more than 50% co-precipitate with CP33A (Teubner et al., 2017), are either no (*psbF*) or are weak (*rbcL*) interaction partners of CP33B. How specificity is generated, remains unclear. Our RBNS analysis uncovered a sequence motif as prime target that is not enriched in *psbA* versus other chloroplast transcripts. It does however prove that CP33B is well capable of binding RNA by itself, without need for auxiliary proteins, which is in line with previous electro mobility shift assays of cpRNPs (Li and Sugiura, 1991; Ye and Sugiura, 1992; Lisitsky et al., 1995). Still, specificity of binding is likely to be modulated by other proteins, as is frequently the case for RRM proteins (Dreyfuss et al., 2002). For example, U2B, a structurally related RBP with two RRMs binds RNA in response to protein-protein interactions (Scherly et al., 1990). Candidates for protein interactions on the *psbA* mRNA are the two RBPs CP33C and SRRP1, which were co-purified in precipitations of the *psbA* mRNA together with CP33B. Speculatively, specificity for *psbA* might be generated by a larger complex of these three or more RBPs.

While at present unclear, how specifically *psbA* is recognized by CP33B, it is remarkable that almost the entire *psbA* mRNA pool is associated with this protein. Given that *psbA* is the most abundant chloroplast mRNA, with an estimated 14.000 molecules per chloroplast (Nakamura et al., 2001), CP33B numbers should at least equal this amount, or, if we assume multiple target sites within the *psbA* mRNA, should be present in excess. *psbA*, like many of the mRNA targets of CP33B encode photosynthetic membrane proteins,. Such mRNAs, when translated, become tethered to membranes via nascent peptide chains being inserted co-translationally into the membrane (Zoschke and Barkan, 2015). CP33B binds within the coding region of *psbA* according to our oligo-RIP-chip results. This is supported by RBNS data, which point to a sequence motif also found within the coding region of *psbA*. Binding to the coding region is likely facilitated by the absence of translating ribosomes that would be expected to remove obstacles like CP33B via their intrinsic helicase activity. Thus, translation would decrease the association of CP33B with *psbA* and this would occur on membranes. This idea is supported by our finding that CP33B prefers stromal *psbA* over membrane-tethered *psbA*. We therefore speculate that CP33B’s main role is centered on ribosome-free, stromal *psbA*. It is however puzzling that mutants of CP33B do not display defects in *psbA* accumulation or translation (Watkins et al., 2019). Redundancy or specific stress conditions might prevent identifying the true function of CP33B for the *psbA* mRNA.

### cpRNPs interact with their target transcripts via multiple binding sites

As mentioned above, cpRNPs interact with many mRNAs, but where exactly does binding take place within a target transcript and can binding motives possibly be identified? In order to answer these questions, the binding sites of CP33B should be described in more detail. In the past, the binding motifs of the splicing factor MatK (Zoschke et al., 2010) and various PPR proteins were deciphered with the help of an oligonucleotide microarray or comparable investigations (oligonucleotide probes on dot-blots of IP fractions) (Schmitz-Linneweber et al., 2006; Pfalz et al., 2009).

However, after performing CP33B RIP-chip analysis and hybridization on an oligonucleotide microarray, no enrichment of isolated short sequence sections occurred, but rather an enrichment of almost all transcript sections of the target RNAs studied (*psbA, ndhB, psaA/psaB/rps14, ndhF* and *psbD/psbC*) was noted. This is in line with the finding that the intact *psbA* transcript can be recovered in CP33B IPs as evidenced by RNA gel blot hybridization (Watkins et al., 2019), whereas in the case of MatK RNA fragments with a length of ~200 nt to ~500 nt and no *trnK* precursor transcripts (~ 2.8 knt) coprecipitated (Zoschke et al., 2010). A similar picture was observed using oligo RIP-chips analysis of CP29A and CP31A from Arabidopsis, as well as *in vitro* approaches for 28RNP from spinach (Lisitsky et al., 1995; Kupsch et al., 2012). 28RNP interacts with part of the coding regions as well as the 3’ and 5’ UTRs of different mRNAs (Lisitsky et al., 1995). Together, these data could indicate that CP33B as well as other cpRNPs have multiple binding sites within their target transcripts.

Such an RNA-association of cpRNPs across multiple binding sites is consistent with their general and global binding behavior, which is in contrast to the specific binding behavior of PPR-proteins. Speculatively, such a multivalent interaction of cpRNPs with their RNA targets is functionally relevant, possibly for RNA protection against degradation, which is a main function of cpRNPs (Nakamura et al., 2001; Kupsch et al., 2012; Teubner et al., 2017).

## Materials and Methods

### Plant material

*Arabidopsis thaliana* cpRNP T-DNA insertion line *cp33b* (SK31607 from the Saskatoon collection) was obtained from the ABRC (Arabidopsis Biological Resource Center) and grown together with wt *A. thaliana* (ecotype Columbia-0) on soil with a 16-h light/8-h dark cycle at 23°C.

### Localization by Fluorescence Microscopy

A full-length cDNA sequence of CP33B was amplified by PCR and cloned via *Xho*I and *Nco*I as a translational fusion with GFP under the control of a 35S promoter. The PEND-dsRed construct under constitutive control of the ubiquitin promoter (pUbi) was kindly provided by Dr R. Lorbiecke and Dr J. Kluth (University of Hamburg, Germany). Mesophyll protoplast preparation and transfection with CP33B:GFP- and PEND-dsRed-fusion constructs was performed as described (Fuss et al., 2013). A Zeiss 510 Meta confocal laser scanning microscope and the ZEISS LSM IMAGE BROWSER software were used for detection and documentation of fluorescence signals.

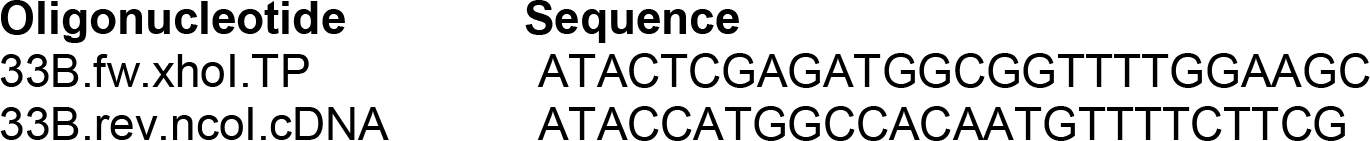

### Dot-Blot Analysis

Dot-blot production was described previously (Teubner et al., 2017). Radiolabeled RNA probes were prepared by *in vitro* transcription of PCR products containing a T7 promoter using T7 RNA polymerase in the presence of α^32^P-UTP.

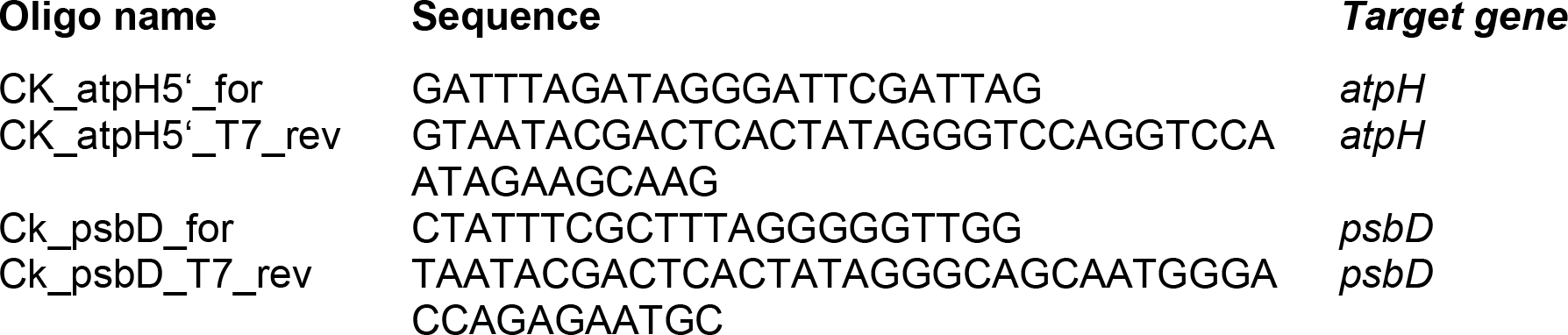
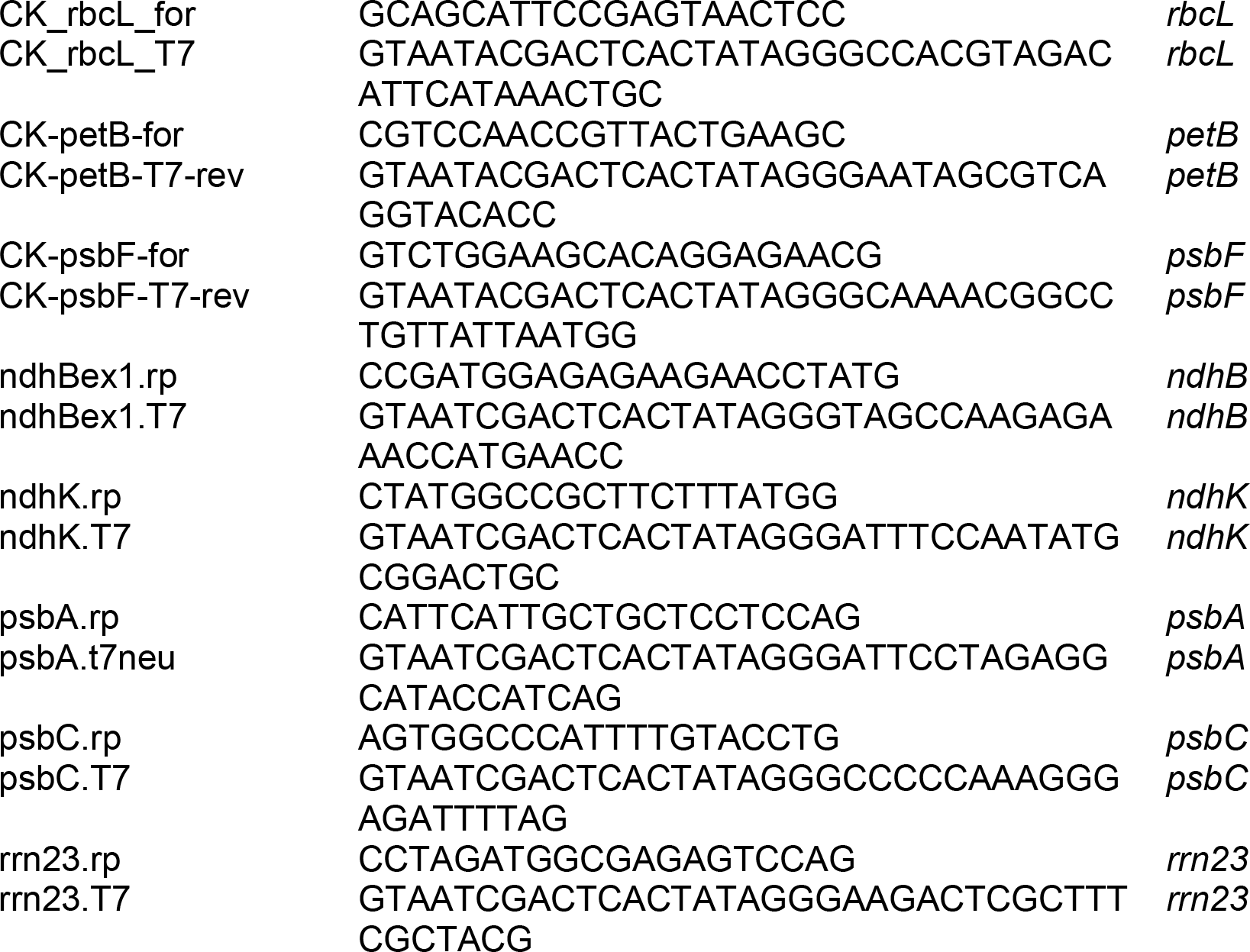

### Immunoblots and antibody production

The production and specificities of the CP33B antisera were reported previously (Watkins et al., 2019). The antisera to PsaD and RuBisCO were obtained from Agrisera. Immunoblots were carried out using standard procedures.

### RIP-chip analysis

CP33B was immunoprecipitated from Arabidopsis chloroplast stroma and co-precipitating RNAs were purified and hybridized to a whole-chloroplast-genome tiling array or a oligonucleotide array, respectively, as previously described (Kupsch et al., 2012). The hybridized microarray were washed, scanned and analyzed as described (Teubner et al., 2017). RIP-chip of CP33B from membrane fractions was performed on membranes prepared from isolated chloroplasts. The membranes were washed five times, solubilized with 1% NP 40 (Nonidet™ P 40) for 15 min on ice and centrifuged (10min, 20,000 × g, 4°C) to remove unsoluble matter. The dissolved membranes were diluted with 1 vol CoIP buffer and incubated with anti CP33B antisera. All further procedures including data evaluation were carried out analogous to that of the Stroma RIP-chip.

### RBNS analysis

#### Cloning of GST:SBP:CP33B

A streptavidin-binding protein (SBP) tag was introduced into the *pGEX-6P-1* vector downstream of the GST tag and the PreScission cleavage site of the vector via *Bam*HI and *Eco*RI restriction sites. The resulting vector was named *pGEX-SBP*. The coding sequence of CP33B was amplified from cDNA without the predicted signal peptide and cloned into the *pGEX-SBP* vector via *Mfe*I and *Xho*I sites using the following oligonucleotides.

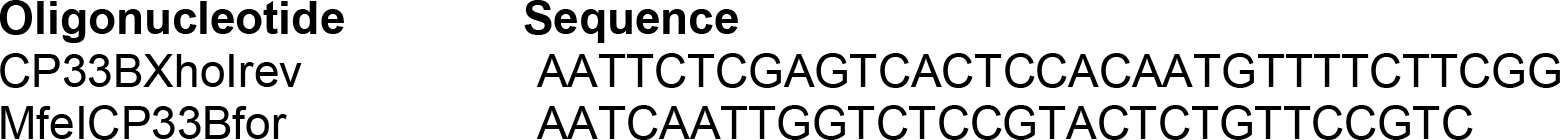

#### RNA oligo production

The RBNS random RNA input for the RNA Bind-n-Seq experiment was prepared by *in vitro* transcription of the RBNS T7 template, a DNA oligo containing a 40mer random sequence flanked by primer sites for the Illumina adapters and a T7 promoter sequence (closely following Lambert et al., 2015). To produce the dsDNA for a more optimal transcription from the T7 polymerase, the T7 promoter oligo was first attached to the RBNS T7 template region corresponding to the T7 promoter. For this purpose, 3 μl of each of the 100 μM oligo samples were heated in water at 65 °C for 5 min and then cooled at room temperature for 2 min. Polymerization of the second DNA strand was performed within 15 minutes at 25 °C in the presence of 33 μM dNTPs, 1× NEB2 buffer and 11 U DNA Polymerase I Large (Klenow) fragment in a 50 μl volume reaction. The reaction was then stopped at 75 °C for 20 min with 10 mM EDTA.

The RBNS Input Oligo Pool was transcribed with T7 RNA Polymerase. DNA was removed using 2 U TURBO™ DNase (Life Technologies) for 5 min at 37 °C. The reaction was stopped by adding 1.5 μl EDTA 0.5 M and incubation at 75 °C for 10 min. The *in vitro* transcribed RNA were purified by gel extraction and concentrated by ethanol precipitation.

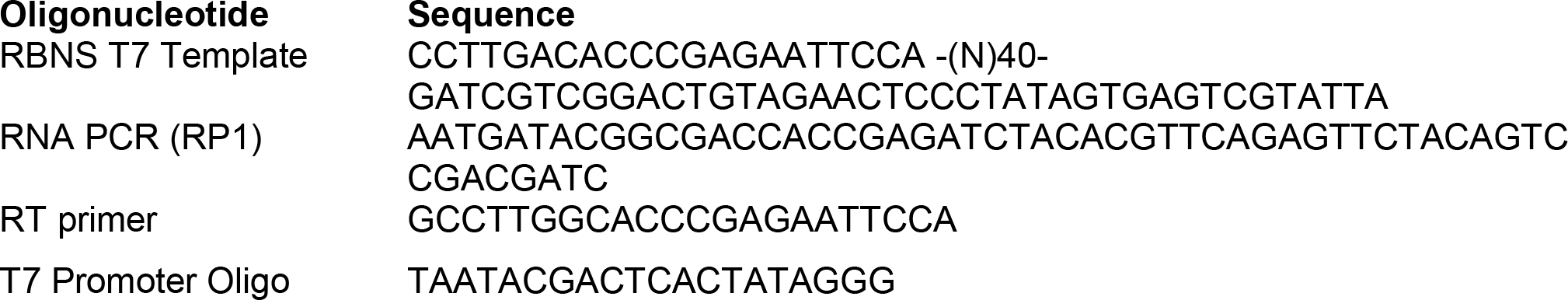

#### RBNS assay

Two different concentrations of SBP-CP33B, 100 nM and 1000 nM were equilibrated in 250 μl binding buffer (25mM Tris-HCl pH 7.5; 150 mM KCl; 3mM MgCl2; 0.01% tween; 1 mg/mL BSA; 1 mM DTT) at 22 °C for 30 min. In addition, a sample with binding buffer only (zero protein sample) was incubated in the same way. The RBNS Input Random RNA was then added to a final concentration of 0.5 μM and incubated for 3 h at 22 °C. Subsequently, streptavidin-coated magnetic beads (Invitrogen) were added and the bead-sample mixture incubated for 1 hour at room temperature. After magnetic separation, the beads were washed with 1 ml Wash Buffer (25mM Tris-HCl pH 7.5; 150 mM KCl; 60 ug/mL BSA; 0.5 mM EDTA; 0.01% tween) and then incubated at 70 °C for 10 min in 100 μl Elution Buffer (10mM Tris-HCl pH 7.0, 1mM EDTA, 1%SDS). RNA from the eluates were purified using the RNA Clean & Concentrator Kit by Zymo Research (Irving, USA). Half of the purified RNA from each protein concentration and the input RNA pool was reverse transcribed using the ProtoScript II reverse transcriptase using the RT primer and subsequently amplified by PCR using the Q5 High-Fidelity DNA polymerase and the RP1 primer (see above) plus barcoded primers. The library was sequenced on a NextSeq 500 (Illumina) at LGC (Berlin). For analysis and motif detection, we used the published bioinformatic pipeline (https://github.com/cburgelab/RBNS_pipeline; Lambert et al., 2014; Dominguez et al., 2018)

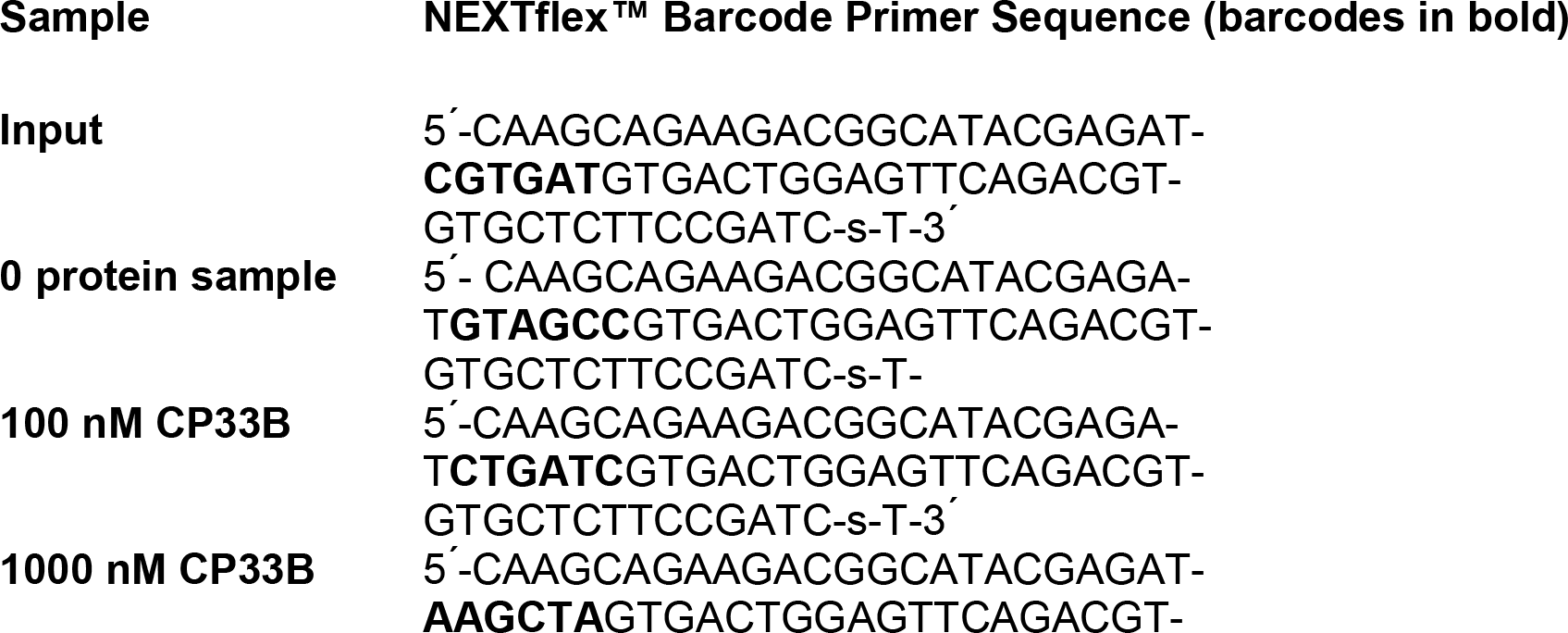

## Acknowledgements

We are grateful to Bernhard Grimm (HU Berlin) for a donation of the GSAT antibody. This work was supported by a grant of the Deutsche Forschungsgemeinschaft within CRC TRR175 to J.N. (project A06) and C.S.L (project A02).

