## Supplemental Table 1 for "The chloroplast ribonucleoprotein CP33B quantitatively binds the *psbA* mRNA"

**Supplemental Table 1**: RIP-Chip data for Fig. 3A

| Name | | | Median(F532 Median - B532) | | Median (Median of Ratios (635/532)) | | Diff. Enrich. [2] |
|---|---|---|---|---|---|---|---|
| | Spot Count CP33B [1] | Spot Count control [1] | CP33B | control | CP33B | control | (CP33B/ control) |
| H-psbA | 36 | 36 | 4896 | 10997 | 0.865 | 0.26964 | 3.207981012 |
| psbA | 36 | 35 | 2968 | 10489 | 19.294 | 0.2604 | 74.093702 |
| psbA-Kex2 | 36 | 36 | 999.5 | 6256.5 | 8.6185 | 0.27636 | 31.18577218 |
| Kex2-matK | 36 | 36 | 775 | 815 | 0.397 | 0.4368 | 0.908882784 |
| matK-1 | 35 | 36 | 507 | 992.5 | 1.449 | 0.59556 | 2.433004231 |
| matK-2 | 35 | 32 | 126 | 297.5 | 0.738 | 0.85596 | 0.862189822 |
| Kint | 30 | 24 | 160 | 2425.5 | 0.8025 | 0.3024 | 2.653769841 |
| Kint-Kex1 | 36 | 25 | 135.5 | 250 | 1.3955 | 0.46704 | 2.987966769 |
| IG_K-rps16 | 35 | 36 | 334 | 877.5 | 1.507 | 0.80724 | 1.866854963 |
| rps16ex2 | 35 | 36 | 629 | 1171 | 1.006 | 0.41748 | 2.409696273 |
| rps16ex2-in-ex1 | 30 | 30 | 173 | 268 | 1.884 | 1.17936 | 1.597476597 |
| rps16in-1 | 27 | 22 | 153 | 2763.5 | 0.57 | 0.10752 | 5.301339286 |
| rps16in-2 | 36 | 30 | 416 | 686 | 0.963 | 0.60144 | 1.601157223 |
| rps16ex2-IG | 27 | 27 | 301 | 341 | 0.538 | 1.28016 | 0.420259968 |
| Q | 36 | 36 | 7156 | 9036.5 | 0.0505 | 0.3024 | 0.166997354 |
| psbK | 36 | 36 | 2897.5 | 6257 | 0.461 | 0.24528 | 1.879484671 |
| psbI-S | 36 | 36 | 1721.5 | 3708.5 | 0.219 | 0.28224 | 0.775935374 |
| IG-psbI-G-1 | 36 | 36 | 18778.5 | 36461 | 0.0445 | 0.21084 | 0.21106052 |
| IG-psbI-G-2 | 36 | 36 | 2310.5 | 3966 | 0.3075 | 0.26292 | 1.16955728 |
| Gex1-in | 36 | 36 | 2531 | 3227.5 | 0.456 | 0.28644 | 1.591956431 |
| in-Gex2 | 36 | 36 | 2946 | 6737 | 0.2665 | 0.25452 | 1.047068993 |
| Gex2-R | 36 | 36 | 1160.5 | 2464.5 | 0.5935 | 0.28056 | 2.115412033 |
| atpA-1 | 35 | 28 | 230 | 320 | 0.61 | 0.80472 | 0.758027637 |
| atpA-3 | 36 | 36 | 355.5 | 831.5 | 1.923 | 0.651 | 2.953917051 |
| atpA-atpFex2 | 36 | 36 | 270.5 | 585 | 2.102 | 0.58716 | 3.579944138 |
| atpFex2-in | 36 | 36 | 857.5 | 2720 | 2.322 | 0.3948 | 5.881458967 |
| atpFex2-in-ex1 | 36 | 36 | 1526.5 | 5452.5 | 1.525 | 0.26712 | 5.709044624 |
| atpFin-ex1 | 36 | 36 | 864 | 2863 | 1.7555 | 0.35952 | 4.882899421 |
| atpFex1-IG | 36 | 36 | 1602 | 4978.5 | 3.223 | 0.30828 | 10.45478137 |
| atpH | 36 | 33 | 239 | 702 | 1.7405 | 0.68544 | 2.539244865 |
| atpH-IG | 36 | 36 | 3138.5 | 6641 | 4.7255 | 0.25452 | 18.56632092 |
| IG-atpI | 32 | 36 | 282 | 1146 | 2.1335 | 0.48132 | 4.432602011 |
| atpI | 36 | 33 | 259.5 | 822 | 3.1145 | 0.672 | 4.634672619 |
| atpI-rps2 | 36 | 36 | 1058 | 3704 | 2.435 | 0.37212 | 6.54358809 |
| rps2 | 36 | 24 | 167.5 | 1488.5 | 0.4105 | 0.56364 | 0.728301753 |
| rps2-IG | 36 | 21 | 81.5 | 5351 | 1.8615 | 0.0672 | 27.70089286 |
| rpoC2-1 | 27 | 24 | 127 | 1139.5 | 1.515 | 0.76776 | 1.973272898 |
| rpoC2-3 | 36 | 27 | 111 | 5268 | 3.1585 | 0.63168 | 5.000158308 |
| rpoC2-4 | 36 | 20 | 131 | 391 | 1.831 | 0.04788 | 38.24143693 |
| rpoC2-5 | 30 | 21 | 146.5 | 6858 | 0.687 | 0.0588 | 11.68367347 |
| rpoC2-6 | 27 | 24 | 129 | 4321 | 1.357 | 0.43176 | 3.142949787 |
| rpoC2-7 | 24 | 24 | 113 | 2622 | 1.1505 | 0.36624 | 3.1413827 |

| | | | | | | | |
|---|---|---|---|---|---|---|---|
| rpoC2-rpoC1-1 | 21 | 25 | 2862 | 7426 | 0.111 | 0.48048 | 0.231018981 |
| rpoC1ex1-1 | 36 | 27 | 3417 | 419 | 1.51 | 0.80976 | 1.864750049 |
| rpoC1ex1-in | 36 | 24 | 1291.5 | 2211 | 1.639 | 0.40488 | 4.048113021 |
| rpoC1ex1-in-ex2 | 36 | 24 | 166.5 | 255 | 1.362 | 0.1848 | 7.37012987 |
| rpoC1in | 36 | 32 | 146 | 2539.5 | 1.869 | 0.91056 | 2.052583026 |
| rpoC1in-ex2 | 29 | 27 | 97.5 | 2744.5 | 2.772 | 0.88872 | 3.119092628 |
| rpoC1-rpoB-1 | 30 | 27 | 148.5 | 2330.5 | 1.5755 | 1.01808 | 1.547520824 |
| rpoC1-rpoB-2 | 36 | 30 | 265 | 378 | 1.064 | 0.6048 | 1.759259259 |
| rpoB-1 | 36 | 36 | 142 | 294 | 1.5725 | 0.91812 | 1.712739076 |
| rpoB-2 | 27 | 23 | 140.5 | 276 | 1.6 | 0.14112 | 11.33786848 |
| rpoB-3 | 30 | 24 | 161.5 | 417 | 0.844 | 0.51576 | 1.63642004 |
| rpoB-4 | 36 | 36 | 211 | 445 | 2.82 | 1.20372 | 2.342737514 |
| rpoB-IG | 24 | 27 | 160 | 3317 | 0.73 | 1.56744 | 0.465727556 |
| IG rpoB-C | 36 | 27 | 94.5 | 3436.5 | 2.3055 | 1.08192 | 2.130933895 |
| IG-C | 36 | 36 | 734.5 | 675.5 | 0.0715 | 0.2814 | 0.254086709 |
| C-IG | 36 | 36 | 1035.5 | 179 | 0.6275 | 0.37464 | 1.674941277 |
| IG-petN | 36 | 36 | 135 | 410 | 0.3795 | 0.27972 | 1.356713857 |
| petN-IG | 36 | 36 | 6955.5 | 11301.5 | 0.6235 | 0.4914 | 1.268823769 |
| psbM | 24 | 20 | 563.5 | 1030 | 0.2075 | 0.08652 | 2.398289413 |
| IG psbM-D-1 | 36 | 36 | 2043.5 | 3477 | 0.413 | 0.37968 | 1.087758112 |
| IG psbM-D-2 | 36 | 34 | 668 | 1012 | 0.7685 | 0.8316 | 0.924122174 |
| D | 36 | 33 | 587.5 | 3066.5 | 0.1735 | 0.32928 | 0.526907191 |
| Y | 35 | 36 | 984.5 | 1112 | 0.575 | 0.42504 | 1.352813853 |
| E | 36 | 36 | 183.5 | 297.5 | 0.4675 | 0.32928 | 1.419764334 |
| IG E-T | 27 | 24 | 1276 | 3006 | 1.345 | 0.75936 | 1.771228403 |
| T | 36 | 36 | 593 | 973.5 | 0.2325 | 0.45276 | 0.513517095 |
| IG T-psbD-1 | 36 | 36 | 2569 | 3560.5 | 1.4555 | 0.48216 | 3.018707483 |
| IG T-psbD-2 | 36 | 36 | 81 | 1691.5 | 1.761 | 0.21924 | 8.032293377 |
| psbD-1 | 36 | 36 | 991 | 1134 | 4.011 | 0.32424 | 12.37046632 |
| psbD-2 | 36 | 36 | 630.5 | 1606.5 | 3.302 | 0.40152 | 8.223749751 |
| psbD-psbC | 36 | 36 | 2249.5 | 5629.5 | 2.3455 | 0.39564 | 5.928369225 |
| psbC-1 | 36 | 36 | 1811.5 | 4857 | 1.6015 | 0.27132 | 5.902624208 |
| psbC-2 | 36 | 36 | 2276.5 | 3569.5 | 2.599 | 0.26292 | 9.885136163 |
| psbC-S | 36 | 36 | 327.5 | 2312 | 1.2265 | 0.27888 | 4.397948939 |
| S-psbZ | 36 | 36 | 2016 | 3570.5 | 2.1935 | 0.33852 | 6.479676238 |
| IG psbZ-G | 36 | 36 | 3267 | 7068 | 0.3125 | 0.31836 | 0.981593165 |
| G-fM | 36 | 36 | 2211.5 | 4579.5 | 0.174 | 0.3696 | 0.470779221 |
| fm-rps14 | 36 | 36 | 1089 | 2343.5 | 0.144 | 0.3024 | 0.476190476 |
| rps14-psaB-1 | 36 | 36 | 1453.5 | 2553 | 1.0915 | 0.28308 | 3.85580048 |
| rps14-psaB-2 | 36 | 36 | 1588 | 1865 | 0.95 | 0.31752 | 2.991937516 |
| psaB-1 | 36 | 36 | 3493 | 4891.5 | 2.3715 | 0.294 | 8.066326531 |
| psaB-2 | 36 | 36 | 1719.5 | 4021 | 1.2615 | 0.41244 | 3.058626709 |
| psaB-psaA-1 | 36 | 36 | 961 | 2540.5 | 1.9165 | 0.58968 | 3.250067833 |
| psaB-psaA-2 | 36 | 27 | 3030 | 5264.5 | 1.588 | 0.32424 | 4.897606711 |
| psaA-1 | 36 | 36 | 422 | 1486 | 1.7435 | 0.31836 | 5.476504586 |
| psaA-2 | 35 | 32 | 333 | 1004 | 1.332 | 0.65268 | 2.040816327 |
| psaA-3 | 36 | 36 | 201 | 2173 | 2.101 | 0.30324 | 6.928505474 |
| psaA-IG-1 | 36 | 36 | 1816.5 | 2713.5 | 1.5685 | 0.31332 | 5.006064088 |
| psaA-IG-2 | 27 | 24 | 307 | 958.5 | 1.992 | 0.61656 | 3.230829116 |
| IG-ycf3ex3 | 36 | 36 | 1206.5 | 3277 | 1.096 | 1.49436 | 0.733424342 |

| | | | | | | | |
|---|---|---|---|---|---|---|---|
| ycf3ex3-in2 | 24 | 33 | 1432.5 | 2560.5 | 1.0835 | 1.23984 | 0.873903084 |
| ycf3ex3-in2-ex2 | 36 | 36 | 137 | 2233 | 0.821 | 0.5628 | 1.458777541 |
| ycf3in2 | 33 | 27 | 187.5 | 242 | 1.977 | 0.80136 | 2.467056005 |
| ycf3ex2 | 36 | 27 | 702 | 296 | 2.2975 | 0.85512 | 2.686757414 |
| ycf3ex1 | 36 | 33 | 342.5 | 577 | 0.7745 | 0.7644 | 1.013212977 |
| S-GGA | 36 | 36 | 202 | 372 | 0.0665 | 0.2478 | 0.268361582 |
| rps4-1 | 32 | 23 | 124 | 473 | 0.9155 | 0.07392 | 12.38501082 |
| rps4-2 | 32 | 33 | 969 | 5234 | 0.979 | 1.04664 | 0.93537415 |
| T-UGU | 35 | 36 | 160.5 | 311 | 0.45 | 0.55524 | 0.810460341 |
| T-IG | 36 | 36 | 3436.5 | 5409 | 0.058 | 0.23184 | 0.250172533 |
| L-UAA ex1-in | 36 | 36 | 161.5 | 4510 | 0.1505 | 0.27888 | 0.539658635 |
| L-UAA in-ex2 | 36 | 36 | 207.5 | 363 | 0.1255 | 0.231 | 0.543290043 |
| L-UAA ex2-IG | 36 | 36 | 512 | 679 | 0.1715 | 0.26376 | 0.650212314 |
| F-GAA-1 | 36 | 36 | 3596 | 5195.5 | 0.0715 | 0.24948 | 0.28659612 |
| F-GAA-2 | 36 | 36 | 8786 | 11786 | 0.159 | 0.29904 | 0.531701445 |
| ndhJ | 36 | 36 | 6748.5 | 10830 | 1.228 | 0.48132 | 2.551317211 |
| ndhJ-ndhK | 36 | 36 | 3352.5 | 6778.5 | 2.1165 | 0.5838 | 3.625385406 |
| ndhK | 36 | 30 | 3395.5 | 6486 | 3.11 | 0.34524 | 9.008226162 |
| ndhK-ndhC | 36 | 36 | 2867 | 4286 | 3.102 | 0.399 | 7.77443609 |
| ndhC-IG | 36 | 36 | 357.5 | 923.5 | 3.4985 | 0.31752 | 11.01820358 |
| IG ndhC-V-UAA | 36 | 24 | 208 | 616.5 | 1.07 | 0.42 | 2.547619048 |
| Vex2-in | 35 | 27 | 217.5 | 2014.5 | 0.684 | 0.23352 | 2.929085303 |
| Vin-ex1 | 36 | 36 | 453 | 1585 | 0.1415 | 0.34944 | 0.404933608 |
| Vex1-atpE | 36 | 36 | 754 | 1863 | 0.0975 | 0.4578 | 0.212975098 |
| atpE-atpB | 35 | 36 | 264 | 3357 | 0.397 | 0.45948 | 0.864020197 |
| atpB-1 | 35 | 36 | 246 | 7685 | 0.465 | 0.48972 | 0.949522176 |
| atpB-2 | 36 | 36 | 1335.5 | 2004 | 0.9215 | 0.71484 | 1.289099659 |
| atpB-3 | 36 | 36 | 2371 | 4049.5 | 0.693 | 0.7644 | 0.906593407 |
| atpB-rbcL | 36 | 36 | 540 | 950.5 | 0.4615 | 0.38556 | 1.196960266 |
| rbcL-1 | 36 | 36 | 1053 | 1727.5 | 1.005 | 0.399 | 2.518796992 |
| rbcL-2 | 36 | 36 | 390.5 | 574.5 | 0.9705 | 0.39816 | 2.437462327 |
| rbcL-3 | 36 | 36 | 892 | 1489 | 1.3665 | 0.44184 | 3.092748506 |
| IG rbcL-accD | 36 | 36 | 1038.5 | 2157.5 | 0.4545 | 0.5124 | 0.887002342 |
| accD-1 | 31 | 34 | 2738.5 | 4672.5 | 0.921 | 0.68796 | 1.338740624 |
| accD-2 | 36 | 36 | 2596 | 4463.5 | 1.122 | 0.61572 | 1.82225687 |
| accD-3 | 36 | 33 | 4543 | 6965 | 0.899 | 0.6468 | 1.389919604 |
| IG accD-psaI | 36 | 36 | 437.5 | 762 | 0.2705 | 0.23856 | 1.133886653 |
| psaI | 36 | 36 | 321 | 332.5 | 1.481 | 0.20328 | 7.285517513 |
| ycf4-1 | 36 | 34 | 366 | 582 | 1.182 | 0.294 | 4.020408163 |
| ycf4-2 | 36 | 33 | 243.5 | 557 | 0.7345 | 0.24696 | 2.974165857 |
| cemA-1 | 36 | 33 | 1148 | 4940.5 | 1.1075 | 0.65688 | 1.686000487 |
| cemA-2 | 36 | 36 | 449.5 | 5280.5 | 0.995 | 0.39228 | 2.536453554 |
| cemA-3 | 36 | 36 | 1268.5 | 9907 | 0.843 | 0.51996 | 1.62127856 |
| petA-1 | 36 | 32 | 296 | 5114 | 0.6795 | 0.2604 | 2.609447005 |
| petA-2 | 36 | 36 | 433 | 553 | 0.701 | 0.546 | 1.283882784 |
| petA-3 | 36 | 36 | 673 | 2189.5 | 0.5895 | 0.59304 | 0.994030757 |
| IG petA-psbJ | 36 | 33 | 910 | 1201.5 | 0.47 | 0.47544 | 0.988557967 |
| psbJLF | 36 | 36 | 275 | 3724 | 0.217 | 0.39816 | 0.545007032 |
| psbFE | 36 | 36 | 1724 | 1813 | 0.3815 | 0.25116 | 1.518952062 |
| IG psbE-petL-1 | 36 | 36 | 1089.5 | 1773.5 | 0.3305 | 0.28056 | 1.178001141 |

| | | | | | | | |
|---|---|---|---|---|---|---|---|
| IG psbE-petL-2 | 36 | 36 | 1063.5 | 1258 | 0.732 | 0.25872 | 2.829313544 |
| IG psbE-petL-3 | 36 | 36 | 6966.5 | 7125 | 0.6015 | 0.44772 | 1.3434736 |
| petL | 36 | 36 | 3610.5 | 5653 | 0.3735 | 0.45444 | 0.821890679 |
| petG-W-CCA | 36 | 36 | 2485 | 3945.5 | 0.2565 | 0.29064 | 0.882535095 |
| P-UGG | 36 | 36 | 434.5 | 3612 | 0.0785 | 0.42084 | 0.186531699 |
| psaJ | 35 | 36 | 525.5 | 910 | 0.113 | 0.38136 | 0.29630795 |
| IG psaJ-rpl33 | 36 | 36 | 1964 | 3208.5 | 0.439 | 0.378 | 1.161375661 |
| rpl33 | 36 | 36 | 1393.5 | 2372 | 0.6425 | 0.63168 | 1.017128926 |
| rpl33-rps18 | 36 | 36 | 5175.5 | 5926 | 0.622 | 0.504 | 1.234126984 |
| rps18 | 35 | 35 | 5103 | 8587 | 0.785 | 0.27888 | 2.814830752 |
| rpl20 | 32 | 32 | 1559 | 2512.5 | 0.5285 | 0.24864 | 2.125563063 |
| rpl20-IG | 33 | 35 | 848 | 1164.5 | 0.652 | 0.40488 | 1.610353685 |
| in1-rps12ex1 | 33 | 36 | 1197.5 | 963 | 0.736 | 0.27132 | 2.712664013 |
| rps12ex1-clpPex3 | 36 | 36 | 349 | 3102 | 0.6255 | 0.38976 | 1.604833744 |
| clpPex3-in2 | 35 | 36 | 207.5 | 4146.5 | 0.464 | 0.73164 | 0.634191679 |
| in2-clpPex2 | 36 | 35 | 224 | 988 | 0.5975 | 0.77448 | 0.771485384 |
| clpPex2-in1 | 36 | 36 | 497 | 2757 | 0.484 | 0.75348 | 0.642352816 |
| clpPin1 | 36 | 36 | 1246.5 | 1671.5 | 0.4755 | 0.36204 | 1.313390786 |
| clpPex1 | 36 | 36 | 606 | 906.5 | 1.173 | 0.24108 | 4.865604778 |
| psbB-1 | 36 | 36 | 316 | 407 | 2.1125 | 0.26964 | 7.834520101 |
| psbB-2 | 36 | 36 | 576.5 | 613.5 | 0.7715 | 0.27804 | 2.774780607 |
| psbBTN | 36 | 36 | 721.5 | 1509.5 | 0.9385 | 0.36372 | 2.580281535 |
| psbNH | 36 | 36 | 913 | 3189.5 | 0.246 | 0.42252 | 0.58222096 |
| psbH-petBex1-in | 36 | 36 | 1915.5 | 4513 | 0.9915 | 0.36792 | 2.694879322 |
| petBin | 36 | 36 | 582 | 2669.5 | 1.6435 | 0.45276 | 3.629958477 |
| petBin-ex2 | 36 | 36 | 597.5 | 4179 | 2.5625 | 0.23604 | 10.85621081 |
| petBex2 | 36 | 36 | 2698.5 | 3685.5 | 1.5335 | 0.52752 | 2.906998787 |
| petBex2-petDex1-in | 36 | 36 | 458.5 | 1613 | 0.9835 | 0.44688 | 2.200814536 |
| petDin | 36 | 36 | 711 | 1521.5 | 0.753 | 0.31668 | 2.377794619 |
| petDex2 | 36 | 36 | 1322.5 | 3166 | 1.73 | 0.30912 | 5.596532091 |
| petD-rpoA | 36 | 36 | 1036.5 | 1403 | 0.473 | 0.43428 | 1.089159068 |
| rpoA | 36 | 36 | 1492 | 2849 | 0.5705 | 0.34944 | 1.632612179 |
| rpoA-rps11 | 33 | 34 | 1268 | 2270.5 | 0.605 | 0.52836 | 1.145052616 |
| rps11-rpl36 | 36 | 24 | 1180 | 1993.5 | 0.948 | 0.14112 | 6.717687075 |
| IG rpl36-rps8 | 36 | 34 | 1583.5 | 2513.5 | 0.8475 | 0.52164 | 1.62468369 |
| rps8 | 33 | 27 | 2664.5 | 4792 | 0.684 | 0.4452 | 1.53638814 |
| rps8-rpl14 | 36 | 36 | 250 | 623 | 0.81 | 0.40908 | 1.980052801 |
| rpl14 | 36 | 36 | 372 | 4212.5 | 1.0635 | 0.40992 | 2.594408665 |
| rpl16ex2 | 36 | 35 | 207.5 | 497.5 | 0.9765 | 0.33264 | 2.935606061 |
| rpl16in-1 | 35 | 36 | 199 | 972 | 0.96 | 0.21252 | 4.517221909 |
| rpl16in-2 | 24 | 24 | 384 | 888 | 0.926 | 0.1428 | 6.484593838 |
| rpl16in-ex1 | 36 | 33 | 503 | 961 | 1.122 | 0.252 | 4.452380952 |
| rps3-1 | 33 | 35 | 350.5 | 7197 | 0.848 | 0.18648 | 4.547404547 |
| rps3-2 | 35 | 33 | 292 | 5128 | 0.937 | 0.2604 | 3.598310292 |
| rpl22 | 36 | 36 | 668.5 | 4204.5 | 1.103 | 0.26124 | 4.222171184 |
| rps19 | 35 | 36 | 195.5 | 6589 | 0.948 | 0.29736 | 3.188054883 |
| rps19-rpl2ex2 | 36 | 36 | 196 | 7091 | 1.9835 | 0.27216 | 7.287992357 |
| rpl2ex2-in | 36 | 36 | 176 | 2227 | 2.017 | 0.24864 | 8.112129987 |
| rpl2ex2-in-ex1 | 36 | 36 | 246.5 | 1590.5 | 3.1065 | 0.48048 | 6.46540959 |

| | | | | | | | |
|---|---|---|---|---|---|---|---|
| rpl2in-ex1 | 36 | 36 | 415 | 1864 | 0.5765 | 0.29904 | 1.927835741 |
| rpl2ex1 | 36 | 36 | 511 | 2374 | 1.733 | 0.39228 | 4.417762822 |
| rpl23-I-CAU | 36 | 34 | 351 | 8055 | 1.796 | 0.9072 | 1.979717813 |
| I-CAU-ycf2 | 36 | 36 | 589 | 1198.5 | 0.138 | 0.41832 | 0.329890993 |
| ycf2-I-CAU | 36 | 36 | 560 | 1020 | 0.176 | 0.3024 | 0.582010582 |
| ycf2-1 | 36 | 26 | 4449.5 | 6352.5 | 0.9375 | 0.86268 | 1.086729726 |
| ycf2-9 | 36 | 28 | 262 | 433 | 1.81 | 0.6132 | 2.951728637 |
| ycf2-3 | 33 | 29 | 429.5 | 1749.5 | 1.238 | 0.75768 | 1.633935171 |
| ycf2-4 | 36 | 30 | 312 | 523.5 | 1.1375 | 0.441 | 2.579365079 |
| ycf2-5 | 34 | 29 | 1879.5 | 2150 | 1.276 | 0.75768 | 1.684088269 |
| ycf2-6 | 36 | 31 | 285 | 321 | 1.9435 | 0.6636 | 2.928722122 |
| ycf2-7 | 36 | 32 | 134 | 275 | 0.9835 | 0.5922 | 1.660756501 |
| ycf2-8 | 36 | 33 | 264.5 | 710 | 0.573 | 0.70224 | 0.815960355 |
| ycf2-IG | 36 | 31 | 216.5 | 345 | 0.698 | 0.25536 | 2.73339599 |
| IG-L-CAA | 36 | 30 | 249.5 | 533 | 0.8115 | 0.30828 | 2.632347217 |
| L-CAA | 33 | 30 | 373.5 | 664 | 0.57 | 0.26712 | 2.133872417 |
| ndhBex2-1 | 36 | 36 | 421 | 477 | 0.76 | 0.40656 | 1.869342778 |
| ndhBex2-2 | 36 | 33 | 209.5 | 9832 | 0.846 | 0.30408 | 2.782162589 |
| ndhBex2-in | 36 | 33 | 306.5 | 1580.5 | 1.226 | 0.2772 | 4.422799423 |
| ndhBex2-in-ex1 | 36 | 36 | 259 | 5917.5 | 1.274 | 0.48636 | 2.619458837 |
| ndhBin-ex1 | 33 | 27 | 373 | 1692.5 | 0.864 | 0.3696 | 2.337662338 |
| ndhBex1-1 | 36 | 36 | 387 | 1136 | 1.405 | 0.62496 | 2.248143881 |
| ndhBex1-2 | 36 | 36 | 275.5 | 6025 | 1.338 | 0.32844 | 4.073803434 |
| IG-rps7 | 36 | 30 | 462.5 | 865.5 | 0.573 | 0.35364 | 1.620291822 |
| rps7-rps12ex3 | 36 | 36 | 241 | 1306 | 0.8185 | 0.30744 | 2.662308093 |
| rps12in2-ex2 | 36 | 36 | 446.5 | 515.5 | 1.649 | 0.70308 | 2.34539455 |
| rps12ex2 | 36 | 36 | 750.5 | 3298.5 | 0.9165 | 0.52332 | 1.751318505 |
| rps12in1 | 36 | 36 | 293.5 | 2545 | 0.892 | 0.42924 | 2.078091511 |
| IG rps12-V-GAC-1 | 36 | 31 | 5859 | 8411 | 3.0565 | 0.90216 | 3.387979959 |
| IG rps12-V-GAC-2 | 35 | 33 | 381 | 933 | 2.3 | 1.0416 | 2.208141321 |
| IG rps12-V-GAC-3 | 36 | 28 | 633.5 | 1296.5 | 2.0415 | 0.85764 | 2.380369386 |
| IG rps12-V-GAC-4 | 33 | 33 | 1222 | 1470 | 1.925 | 0.82656 | 2.328929539 |
| IG rps12-V-GAC-5 | 35 | 29 | 158.5 | 483 | 1.553 | 0.72912 | 2.129964889 |
| V-GAC | 36 | 36 | 232 | 359 | 0.3415 | 0.32928 | 1.037111273 |
| V-GAC-16S | 36 | 36 | 172 | 492.5 | 0.098 | 0.30996 | 0.316169828 |
| 16S-1 | 36 | 36 | 503 | 445 | 0.0925 | 0.28896 | 0.320113511 |
| 16S-2 | 36 | 36 | 265 | 585 | 0.0955 | 0.2898 | 0.329537612 |
| 16S-3 | 36 | 36 | 1464 | 3246.5 | 0.083 | 0.31584 | 0.262791287 |
| 16S-4 | 36 | 36 | 23984.5 | 30137 | 0.0785 | 0.26796 | 0.292954172 |
| 16S-5 | 36 | 36 | 27257 | 48301 | 0.2145 | 0.27552 | 0.778527875 |
| I-GAU ex1 | 36 | 36 | 15318.5 | 35822.5 | 0.62 | 0.2898 | 2.139406487 |
| I-GAU ex1-in | 24 | 36 | 24784.5 | 28595 | 3.1935 | 0.357 | 8.945378151 |
| I-GAU ex2 | 36 | 36 | 21282.5 | 34325.5 | 0.97 | 0.28056 | 3.457370972 |
| I-GAU-A-UGC | 36 | 36 | 10051.5 | 20862.5 | 0.314 | 0.3696 | 0.8495671 |
| A-UGC ex1-in | 36 | 36 | 8104.5 | 14494 | 0.574 | 0.36792 | 1.560121766 |
| A-UGC -in | 36 | 36 | 2763.5 | 6826 | 0.588 | 0.41412 | 1.419878296 |
| A-UGC ex1 | 36 | 36 | 7812.5 | 9685 | 0.451 | 0.32424 | 1.390944979 |
| A-UGC-23S | 36 | 36 | 2096 | 4109.5 | 0.1625 | 0.22008 | 0.738367866 |
| 23S-1 | 36 | 36 | 1198.5 | 3319.5 | 0.168 | 0.20328 | 0.826446281 |
| 23S-2 | 36 | 36 | 2340 | 3639 | 0.1475 | 0.22092 | 0.667662502 |

| | | | | | | | |
|---|---|---|---|---|---|---|---|
| 23S-3 | 36 | 36 | 2332.5 | 4745 | 0.177 | 0.23856 | 0.74195171 |
| 23S-4 | 36 | 36 | 23287 | 39978 | 0.191 | 0.20244 | 0.943489429 |
| 23S-5 | 36 | 36 | 19590 | 36516 | 0.1565 | 0.23016 | 0.679961766 |
| 23S-6 | 36 | 36 | 7680.5 | 21798 | 0.1625 | 0.25284 | 0.64269894 |
| 23S-7 | 36 | 36 | 24805.5 | 41495 | 0.175 | 0.24024 | 0.728438228 |
| 23S-8 | 36 | 36 | 35717 | 45359 | 0.1925 | 0.23604 | 0.815539739 |
| 23S-9 | 36 | 36 | 20270 | 36888 | 0.0575 | 0.20328 | 0.282861078 |
| 4.5S | 36 | 36 | 12367 | 25611.5 | 0.0785 | 0.23688 | 0.331391422 |
| R-ACG-1 | 36 | 36 | 29521.5 | 43793 | 0.283 | 0.27972 | 1.011726012 |
| R-ACG-2 | 36 | 35 | 13801 | 13724.5 | 1.711 | 0.33768 | 5.066927268 |
| N-GUU-1 | 35 | 36 | 20210.5 | 31881.5 | 0.634 | 0.30072 | 2.108273477 |
| IG-ycf1 | 36 | 36 | 12483 | 22290 | 0.17 | 0.3108 | 0.546975547 |
| 3'ndhF(ycf1) | 36 | 36 | 5159 | 8611.5 | 1.4305 | 0.50316 | 2.843032038 |
| ndhF-1 | 36 | 36 | 364 | 2107 | 1.1915 | 0.315 | 3.782539683 |
| ndhF-2 | 36 | 36 | 754 | 4614 | 0.8035 | 0.35868 | 2.240158358 |
| ndhF-3 | 36 | 36 | 3847.5 | 6334.5 | 1.19 | 0.32172 | 3.698868581 |
| ndhF-4 | 36 | 36 | 1211 | 2107.5 | 2.312 | 0.54012 | 4.280530253 |
| ndhF-5 | 36 | 36 | 740.5 | 1149 | 0.9015 | 0.27972 | 3.222865723 |
| ndhF-IG | 36 | 36 | 997 | 1381.5 | 0.5955 | 0.32004 | 1.860704912 |
| IG ndhF-rpl32 | 33 | 30 | 637 | 1399.5 | 0.341 | 0.3192 | 1.068295739 |
| rpl32 | 36 | 33 | 497 | 1010 | 0.1315 | 0.19656 | 0.669006919 |
| IG rpl32-ccsA | 27 | 20 | 4689.5 | 9247.5 | 1.933 | 0.04368 | 44.253663 |
| IG-ccsA | 33 | 33 | 365.5 | 1137.5 | 0.75 | 0.40488 | 1.852400711 |
| ccsA | 36 | 36 | 210 | 539.5 | 3.9475 | 0.35616 | 11.0835018 |
| ccsA-ndhD | 33 | 24 | 1031 | 1707 | 2.977 | 0.26544 | 11.21534057 |
| ndhD-1 | 36 | 36 | 90 | 5625 | 1.1665 | 0.48216 | 2.419321387 |
| ndhD-2 | 36 | 26 | 280 | 530 | 1.4445 | 0.54096 | 2.670252884 |
| ndhD-psaC | 36 | 36 | 1045 | 2450.5 | 1.625 | 0.32424 | 5.011719714 |
| psaC-ndhE | 36 | 36 | 101 | 3188 | 0.575 | 0.26544 | 2.166214587 |
| ndhE | 24 | 36 | 579.5 | 820.5 | 1.537 | 0.30576 | 5.02681842 |
| IG-ndhG | 36 | 36 | 609.5 | 833.5 | 0.5445 | 0.23772 | 2.290509844 |
| ndhG | 36 | 36 | 1153 | 3314 | 0.977 | 0.3612 | 2.704872647 |
| ndhG-ndhI | 36 | 36 | 4404 | 10496 | 0.792 | 0.2352 | 3.367346939 |
| ndhI | 36 | 36 | 1457.5 | 1946.5 | 1.4315 | 0.44436 | 3.221487083 |
| ndhI-ndhAex2 | 36 | 36 | 1628 | 3493 | 1.6075 | 0.53676 | 2.994820777 |
| ndhAex2-in | 36 | 36 | 1328.5 | 2354 | 1.7155 | 0.46032 | 3.726755301 |
| ndhAex2-in-ex1 | 36 | 35 | 1880 | 4248.5 | 0.6435 | 0.4452 | 1.44541779 |
| ndhAin-1 | 36 | 34 | 545 | 852 | 0.8755 | 0.48972 | 1.787756269 |
| ndhAin-2 | 36 | 36 | 335 | 877.5 | 0.819 | 0.47208 | 1.734875445 |
| ndhAin-ex1 | 36 | 36 | 593 | 772.5 | 1.6985 | 0.54096 | 3.139788524 |
| ndhAex1-ndhH | 36 | 36 | 1058.5 | 1376 | 1.389 | 0.39564 | 3.510767364 |
| ndhH-1 | 36 | 26 | 503 | 965 | 2.7435 | 0.66528 | 4.123827561 |
| ndhH-2 | 36 | 33 | 438 | 803.5 | 2.718 | 0.95928 | 2.833375031 |
| rps15 | 36 | 33 | 412 | 832 | 2.064 | 0.93744 | 2.201740911 |
| rps15-IG | 36 | 27 | 892 | 1676 | 3.989 | 0.96096 | 4.151057276 |
| ycf1-1 | 36 | 36 | 147 | 346 | 0.713 | 0.28056 | 2.54134588 |
| ycf1-2 | 36 | 36 | 136.5 | 547 | 0.555 | 0.29568 | 1.877029221 |
| ycf1-3 | 36 | 33 | 211.5 | 365 | 1.848 | 0.67872 | 2.722772277 |
| ycf1-4 | 33 | 30 | 103.5 | 371 | 2.451 | 1.12812 | 2.172641208 |
| ycf1-5 | 35 | 30 | 3516.5 | 9785 | 2.7 | 1.27932 | 2.110496201 |

| | | | | | | | |
|---|---|---|---|---|---|---|---|
| ycf1-6 | 36 | 31 | 2938 | 28890.5 | 3.075 | 1.18104 | 2.603637472 |
| ycf1-7 | 36 | 36 | 155.5 | 365 | 1.1995 | 0.60396 | 1.986058679 |
| ycf1-8 | 36 | 31 | 142 | 330.5 | 1.1665 | 0.59808 | 1.950407972 |
| ycf1-9 | 36 | 36 | 94 | 161.5 | 0.8695 | 0.4746 | 1.832069111 |
| N-GUU-2 | 36 | 36 | 98.5 | 239 | 0.1785 | 0.3066 | 0.582191781 |
| IG N-R | 36 | 36 | 474 | 705.5 | 1.072 | 0.73584 | 1.456838443 |

[1] each probe is spotted with 12 replications per chip; spots with less than 5 detectable spots were removed from the d

[2] differential enrichment of IP over control- IP, based on the median of ratios of red (635 nm) or green (532 nm) fluore

lata set
scence
