## Supplemental Table 2 for "The chloroplast ribonucleoprotein CP33B quantitatively binds the *psbA* mRNA"

**Supplemental Table 2**: Oligo RIP-Chip data for Fig. 3B and Fig. S1

| Name | sequence of oligonucleotide | length of probe (nt) | start of probe[1] | end of probe[1] | Spot Count CP33B [2] | Spot Count control [2] | Median(F635 Median - B635) CP33B [3] | Median(F635 Median - B635) control [3] | Median(F532 Median - B532) CP33B [3] | Median(F532 Median - B532) control [3] | Median(Median of Ratios (635/532)) CP33B [4] | Median(Median of Ratios (635/532)) control [4] | Diff. Enrich. [5] |
|---|---|---|---|---|---|---|---|---|---|---|---|---|---|
| psbA-1 | taggaatatttttttccaaatcatatgaatacgattgaatacgatccgtag | 50 | 1667 | 1716 | 48 | 27 | 68 | 9 | 79.5 | 90 | 1.27 | 0.20 | 6.53 |
| psbA-2 | atagccatgtcaaccaatgtaaaatggataagatccttttagtttagatt | 50 | 1547 | 1596 | 36 | 33 | 59.5 | 14 | 89.5 | 85 | 0.86 | 0.28 | 3.13 |
| psbA-3 | gagcttgttatgaaacagtataacatgacttatatagccatgtcaaccaa | 50 | 1514 | 1563 | 48 | 33 | 111 | 25 | 96 | 94 | 1.69 | 0.20 | 8.31 |
| PsbA-4 | gcgcacaaattctctaagtagataattgagagcttgttatgaaacagtat | 50 | 1485 | 1534 | 48 | 30 | 40394 | 1313.5 | 7315.5 | 25633.5 | 5.64 | 0.06 | 90.95 |
| psbA-5 | tttatttaataatcagggtataactcccaagcgcacaaattctctaagta | 50 | 1459 | 1504 | 48 | 33 | 29363 | 1413 | 7950.5 | 27299 | 4.85 | 0.06 | 86.57 |
| psbA-6 | ctaaaattgcagtcatggtaaaatccttggtttatttaataatcagggta | 50 | 1429 | 1476 | 48 | 33 | 28292.5 | 2104 | 5956 | 28785 | 6.53 | 0.07 | 98.98 |
| psbA-7 | aagcgaccccataggctttcgctttcgcgtctctctaaaattgcagtcat | 50 | 1395 | 1444 | 48 | 30 | 44462.5 | 2041 | 6804.5 | 35430.5 | 7.46 | 0.06 | 124.41 |
| psbA-8 | taaggtagggatcatcaaaacaccaaaccatccaatgtaaagacggtttt | 50 | 1322 | 1371 | 48 | 33 | 39141.5 | 1153 | 5379 | 19246 | 6.86 | 0.07 | 102.40 |
| psbA-8a | actggaggagcagcaatgaatgcgataataaaaacagaagttgcggtcaa | 50 | 1272 | 1321 | 24 | 12 | 14389.5 | 1325.5 | 5979 | 15505 | 2.91 | 0.09 | 33.25 |
| psbA-9 | cacgaataccatcaatatctactggaggagcagcaatgaatgcgataata | 50 | 1252 | 1301 | 48 | 33 | 32427.5 | 2353 | 7123 | 30672 | 4.81 | 0.10 | 50.11 |
| psbA-9a | ggcaccggaaataatattgtttccgtaaagaagagatccagaaacaggtt | 50 | 1202 | 1251 | 24 | 12 | 7463.5 | 1546 | 7606 | 18948 | 1.89 | 0.08 | 22.69 |
| psbA-9b | tcccagattgggtaaaaatgcaatccaatagctgcagaagtaggaataat | 50 | 1152 | 1201 | 24 | 12 | 19085 | 1642 | 6884.5 | 21050 | 3.87 | 0.08 | 47.25 |
| psbA-9c | ttagttcataaggaccgccgttgtatagccattcatcaacggatgcagct | 50 | 1102 | 1151 | 24 | 12 | 11171 | 2530.5 | 7021 | 27748.5 | 7.14 | 0.09 | 75.93 |
| psbA-9d | ccactcacgacccatataacaagctacaccaagtaaaaagtgtagaacaa | 50 | 1052 | 1101 | 24 | 12 | 24155.5 | 2220.5 | 10961.5 | 25620 | 4.43 | 0.09 | 49.45 |
| psbA-10 | aggacgcatacccagacggaaactaagttcccactcacgacccatataac | 50 | 1022 | 1071 | 48 | 30 | 30714 | 1757 | 3589 | 28843 | 12.85 | 0.07 | 191.86 |
| psbA-10a | aaaacagcagtcgcagctgcaacaggagctgaatatgcaacagcaatcca | 50 | 972 | 1021 | 24 | 12 | 9143 | 1447.5 | 4228 | 16924 | 3.02 | 0.09 | 33.88 |
| psbA-10b | ctagaggcataccatcagaaaaacttccctgaccaattggatagatcaag | 50 | 922 | 971 | 24 | 12 | 25687 | 2615.5 | 7574 | 26056.5 | 4.73 | 0.10 | 47.58 |
| psbA-10c | gttgtgctcagcctggaatacaatcataaagttgaaagtaccagagattc | 50 | 872 | 921 | 24 | 12 | 16756.5 | 1650.5 | 6521 | 20020 | 5.80 | 0.08 | 70.67 |
| psbA-10d | ccgccgaatacaccagctacacctaacatgtgaaatgggtgcataagaat | 50 | 822 | 871 | 24 | 12 | 21139.5 | 1975 | 6143 | 23275 | 5.58 | 0.09 | 64.11 |
| psbA-10e | tgatcaaactagaagttaccaaggaaccatgcatagcactaaaaagggag | 50 | 772 | 821 | 24 | 12 | 17021.5 | 2277 | 6088.5 | 23926.5 | 4.66 | 0.10 | 45.94 |
| psbA-11 | cattagcagattcattttctgtggtttccctgatcaaactagaagttacc | 50 | 742 | 791 | 48 | 33 | 29557.5 | 1105 | 5638.5 | 21516 | 6.57 | 0.06 | 111.32 |
| psbA-11a | agcagctacaatgttgtaagtttcttcttcttgcccgaatctgtaacctt | 50 | 692 | 741 | 24 | 12 | 35475 | 5512.5 | 8766 | 53834.5 | 7.66 | 0.11 | 68.38 |
| psbA-11b | gaattgttgaaactagcatattggaaaatcaatcggccaaaataaccgtg | 50 | 642 | 691 | 24 | 12 | 5377 | 2866 | 3421.5 | 29905.5 | 13.38 | 0.10 | 140.81 |
| psbA-11c | accaaatacctactaccggccaagccgctaagaagaaatgtaaagaacga | 50 | 592 | 641 | 24 | 12 | 33079.5 | 3175 | 3191 | 31380 | 16.73 | 0.10 | 166.50 |
| psbA-11d | attgaaaccatttaggttgaaagccatagtactaatacctaaagcagtaa | 50 | 542 | 591 | 24 | 12 | 27589 | 3315.5 | 3255.5 | 34282.5 | 11.23 | 0.10 | 111.14 |
| psbA-12 | tccttgactatcaactactgattggttgaaattgaaaccatttaggttga | 50 | 512 | 561 | 48 | 33 | 27935.5 | 1120 | 2757.5 | 19581 | 12.11 | 0.06 | 201.89 |
| psbA-12a | cggttaataatatcagcccaagtattaataacacgtccttgactatcaac | 50 | 477 | 526 | 24 | 12 | 30931.5 | 3680.5 | 3929.5 | 40838 | 18.35 | 0.09 | 199.47 |
| psbA-13 | tacgttcatgcataacttccataccaaggttagcacggttaataatatca | 50 | 442 | 491 | 48 | 30 | 46157 | 1372.5 | 2605 | 32042 | 13.87 | 0.06 | 247.68 |
| psbA-13a | aacagcagctaggtctagagggaagttgtgagcattacgttcatgcataa407 | 53 | 407 | 456 | 24 | 12 | 7107.5 | 1765 | 3836.5 | 19696 | 13.95 | 0.09 | 156.70 |

| | | | | | | | | | | | | | |
|---|---|---|---|---|---|---|---|---|---|---|---|---|---|
| psbA-14 | acactaacgaattatccatttgtagatggagcctcaacagcagctaggtc | 50 | 372 | 421 | 48 | 33 | 32304.5 | 1594 | 2694.5 | 26739 | 13.85 | 0.06 | 216.40 |
| psbA-15 | aagaaggcttatattgctcgtttttttactaaactagatctagactaacac | 50 | 326 | 375 | 48 | 30 | 26378 | 490.5 | 3812.5 | 12585 | 5.88 | 0.06 | 105.92 |
| psbA-16 | acgagtaatataagccctctttcttatttaaagaaggcttatattgctcg | 50 | 296 | 345 | 48 | 33 | 670.5 | 32 | 203 | 461 | 5.31 | 0.07 | 71.76 |
| psbA-17 | ataaaaatttgctcatttttatagaaaaaaacgagtaatataagccctct | 50 | 266 | 315 | 48 | 33 | 1204.5 | 48 | 310.5 | 730 | 4.40 | 0.06 | 71.02 |
| psbA-17a | ttattattatttattattaatataataaataaagtaaaatatgatactct | 50 | 216 | 265 | 23 | 12 | 697 | 27 | 141 | 414 | 4.92 | 0.08 | 58.92 |
| psbA-18a | tagcaaattccaccttattttttttctaataaaaaatatatagtaatttt | 50 | 156 | 205 | 23 | 12 | 89 | 20 | 120 | 281 | 0.81 | 0.09 | 8.71 |
| psbA-19 | tgaacccgcgatggtgaattcacaatccactgccttaatccacttggcta | 50 | 20 | 69 | 48 | 33 | 1420.5 | 1570 | 31671 | 27725 | 0.07 | 0.06 | 1.16 |
| rps14 | ttttccaatttttgcctcttcttctccctataaatcaaacttttctttgc | 50 | 37188 | 37237 | 48 | 30 | 517 | 66.5 | 884 | 777.5 | 0.883 | 0.0685 | 12.89051095 |
| rsp14-psaB | aataaatataagagtgcatacctatattttttttattaaaaaaaatatat | 50 | 37300 | 37349 | 48 | 30 | 120 | 18.5 | 174 | 153.5 | 0.6735 | 0.1215 | 5.543209877 |
| psaB1 | ttaaccgaatttgcccgatgtggaggcaatcaagaaagccgcataagtga | 50 | 37375 | 37424 | 48 | 30 | 226.5 | 38.5 | 408 | 530.5 | 0.7795 | 0.0785 | 9.929936306 |
| psaB2 | atatccattgataagttgtgaagagtttaaccatagataatctcttaacc | 50 | 37675 | 37724 | 48 | 30 | 484.5 | 73 | 959.5 | 892.5 | 0.6185 | 0.0815 | 7.588957055 |
| psaB3 | tactaagatcaatgtagttgtatgtaaacctaaagcaatagcatgatgaa | 50 | 37975 | 38024 | 48 | 30 | 408.5 | 67 | 731.5 | 798 | 0.786 | 0.0795 | 9.886792453 |
| psaB4 | tcccaaagtatggaaccctagaaagaggctggcccaacttaaatgggata | 50 | 38275 | 38324 | 48 | 30 | 124 | 27.5 | 254.5 | 287 | 0.6015 | 0.1045 | 5.755980861 |
| psaB5 | gcctaattgaaaatgaatcgaattattgattgtgtcataaagacccttat | 50 | 38575 | 38624 | 48 | 30 | 402 | 64.5 | 759.5 | 964 | 0.5465 | 0.068 | 8.036764706 |
| psaB6 | ttgagcataaagattccactgacccgtaaaaagtgggcctaacccttggg | 50 | 38875 | 38924 | 48 | 29 | 343.5 | 59 | 767.5 | 856 | 0.5465 | 0.07 | 7.807142857 |
| psaB7 | aagatcttcattagtacgtaaaccgattgtataccaccactgataaacac | 50 | 39175 | 39224 | 48 | 30 | 404.5 | 60 | 763 | 802.5 | 0.739 | 0.08 | 9.2375 |
| psaB8 | atcatgactctcgaagtcatgtgcggtagcaataccaaaccaaatacgac | 50 | 39475 | 39524 | 48 | 30 | 272.5 | 45 | 378.5 | 487 | 0.8335 | 0.092 | 9.059782609 |
| psaB-psaA | ttggctaaaccttggaaatcttaatgccataatgcctttcaaatcctcc | 49 | 39550 | 39598 | 48 | 30 | 484.5 | 72.5 | 694 | 744 | 0.872 | 0.0725 | 12.02758621 |
| psaA1 | tagccattatcctactgcaataattcttgctaagaagaacgcccatgtt | 49 | 39605 | 39647 | 48 | 30 | 422.5 | 69.5 | 723.5 | 742 | 0.7775 | 0.0775 | 10.03225806 |
| psaA2 | tgcccataagaaatcgcggagccacccattaatagtaatggaactctgtg | 50 | 39905 | 39954 | 36 | 24 | 311.5 | 53.5 | 416 | 467.5 | 0.753 | 0.124 | 6.072580645 |
| psaA3 | acctttcaacagtatcaataccgtcacatgaattgtaaatgcatgaatat | 50 | 40205 | 40254 | 48 | 36 | 505 | 75.5 | 867 | 692 | 0.7415 | 0.0945 | 7.846560847 |
| psaA4 | accaaaactgtggaagcctagaaatatacatacccagttgagatgtgata | 50 | 40505 | 40554 | 48 | 36 | 341 | 60 | 640.5 | 484.5 | 0.828 | 0.104 | 7.961538462 |
| psaA5 | agataattgagcatgccatgatgttgttagaatttcatataggcctttat | 50 | 40805 | 40854 | 48 | 34 | 299.5 | 49 | 629 | 469 | 0.5715 | 0.0955 | 5.984293194 |
| psaA6 | ataaagttgagccaaaagatcccgattcaagataaattcatgaggaagcg | 50 | 41105 | 41154 | 48 | 33 | 177.5 | 41 | 368.5 | 294 | 0.5935 | 0.095 | 6.247368421 |
| psaA7 | agttattccggatgctcgccaaatctgaaaaaagcctgaggttatttgta | 50 | 41405 | 41454 | 48 | 33 | 211.5 | 39 | 412 | 396 | 0.626 | 0.117 | 5.35042735 |
| psaA8 | agcatgtaggttccagatccaagtggtagtatcaggtcccttagctattg | 50 | 41705 | 41754 | 48 | 33 | 393 | 65 | 693.5 | 717 | 0.6375 | 0.09 | 7.083333333 |
| psaA-IG | gaataatcattgagtcctcctctttccggacaacacatacaaagaaacc | 49 | 41848 | 41896 | 48 | 33 | 315.5 | 61 | 749 | 678 | 0.596 | 0.089 | 6.696629213 |
| psaA-410 | attatcctatttcaataaatgcttattagtcattactaagaaaaagtcta | 50 | 42217 | 42266 | 40 | 28 | 49 | 17 | 63.5 | 57.5 | 0.7545 | 0.4345 | 1.736478711 |
| ndhB-1 | aaaaaaaccaatttttgatttttggaatggaatatttacggaatccccat | 50 | 97241 | 97192 | 48 | 33 | 71.5 | 21 | 323 | 249 | 0.362 | 0.089 | 4.06741573 |
| ndhB-2 | attctcgtagaatcgagaatgaaattttcattctgtacatgccagatcat | 50 | 97164 | 97115 | 48 | 33 | 73.5 | 18 | 247.5 | 187 | 0.442 | 0.158 | 2.797468354 |
| ndhB-3 | atcggaacaatagggccgttatgctcattacgaaacttgttgaagagatg | 50 | 96979 | 96930 | 48 | 32 | 83.5 | 20 | 251 | 170 | 0.5355 | 0.147 | 3.642857143 |
| ndhB-4 | ttcctcctagagtagctgttaatatgaataacagaaactctgttatagcc | 50 | 96793 | 96744 | 48 | 33 | 66.5 | 26 | 238 | 186 | 0.3645 | 0.122 | 2.987704918 |
| ndhB-5 | tagccaagagaaaccatgaaccagaatagaagagcttgccccacccatga | 50 | 96611 | 96562 | 48 | 30 | 94 | 19.5 | 332 | 326.5 | 0.323 | 0.0845 | 3.822485207 |
| ndhB-6 | tccttcgtatacgtcaggagtccattgatgagaaggggctagggaaagct | 50 | 96437 | 96388 | 48 | 30 | 31.5 | 10.5 | 117 | 94 | 0.425 | 0.1775 | 2.394366197 |
| ndhB-7 | gaaaagcaacgactggagtgggagatccttcgtatacgtcaggagtccat | | | | 48 | 30 | 24 | 9.5 | 96 | 86 | 0.3815 | 0.2065 | 1.847457627 |

| | | | | | | | | | | | | | |
|---|---|---|---|---|---|---|---|---|---|---|---|---|---|
| ndhB-8 | tagagaggtaggaatttctcaaacgaaccgcactccttcgtatacgtcag | 50 | 96404 | 96355 | 45 | 30 | 32 | 11 | 124 | 92.5 | 0.393 | 0.199 | 1.974874372 |
| ndhB-9 | gcatgtccatagagttttgaaaaatccaaacatctcagagatagatagag | 50 | 96359 | 96310 | 44 | 30 | 27 | 13.5 | 116.5 | 90 | 0.5025 | 0.1685 | 2.982195846 |
| ndhB-10 | gtcaaagttctgtcttggtctgcgtggggatagcatttctcttctgcatg | 50 | 96314 | 96265 | 48 | 30 | 82.5 | 16.5 | 127 | 104.5 | 0.6805 | 0.162 | 4.200617284 |
| ndhB-11 | ttcgttgttcctgaccctgcttcaccttaattgttatttgaacaagtcaa | 50 | 96269 | 96220 | 48 | 30 | 49 | 11.5 | 160 | 143.5 | 0.495 | 0.1195 | 4.142259414 |
| ndhB-12 | ccgtaataacaaacttgcaaaatggatccgtttatcataaagagattcgt | 50 | 96224 | 96175 | 46 | 30 | 35 | 12.5 | 119 | 86.5 | 0.674 | 0.2225 | 3.029213483 |
| ndhB-13 | gtctttcttgtatacgtcattagtccgatcctttgcaggaactacccgta | 50 | 96179 | 96130 | 47 | 30 | 49 | 16.5 | 139 | 129 | 0.542 | 0.132 | 4.106060606 |
| ndhB-14 | tgaaggatgagaaccaactatgtagcatctacatcgagaattcaagtctt | 50 | 96134 | 96085 | 47 | 30 | 47 | 17.5 | 139 | 152 | 0.54 | 0.154 | 3.506493506 |
| ndhB-15 | ggtgatccttttgtcgacggatgctcctattacactcgtagtctctgaag | 50 | 96089 | 96040 | 47 | 30 | 33 | 15.5 | 132 | 122.5 | 0.469 | 0.1415 | 3.314487633 |
| ndhB-16 | atctgatttgattcgttctcaatagccatgagattatcatcttagggtga | 50 | 96044 | 95995 | 48 | 30 | 45.5 | 16 | 157 | 139 | 0.4995 | 0.1405 | 3.555160142 |
| ndhB-17 | ccttggttccgtaggagcacgtccgaaagattgagaaatggaaccatctg | 50 | 95999 | 95950 | 48 | 33 | 63.5 | 33 | 193 | 141 | 0.4605 | 0.442 | 1.041855204 |
| ndhB-18 | cttcatcagtggttgtgaatgactgatttttctcaatcttttcgaccttg | 50 | 95954 | 95905 | 48 | 33 | 46.5 | 17 | 132 | 113 | 0.4905 | 0.154 | 3.185064935 |
| ndhB-19 | tgtataagatcgaatcctttctatttctaaaacggattactaatccttaa | 50 | 95890 | 95841 | 48 | 33 | 30.5 | 15 | 105 | 85 | 0.4795 | 0.219 | 2.189497717 |
| ndhB-20 | agtgataaaagtaaagaagaactcatcttctttctttttgattactttc | 50 | 95832 | 95783 | 39 | 33 | 40 | 8 | 119 | 82 | 0.34 | 0.206 | 1.650485437 |
| ndhB-21 | caaaaccgtgcatgagactttcatctcgcacggctcctaagtgataaaag | 50 | 95793 | 95744 | 48 | 33 | 28.5 | 10 | 98 | 88 | 0.4345 | 0.254 | 1.710629921 |
| ndhB-22 | gtcagagtcgaaaagaggattcctcacttctttctctcattcaaaaccgt | 50 | 95752 | 95703 | 48 | 33 | 38 | 17 | 134.5 | 129 | 0.497 | 0.155 | 3.206451613 |
| ndhB-23 | tccgatttgacctatggacgaatatgcaagcatacgtttcatgcttgttt | 50 | 95530 | 95481 | 48 | 33 | 92.5 | 29 | 350 | 341 | 0.3725 | 0.09 | 4.138888889 |
| ndhB-24 | gagctaaagagagagccaaaaaaggatcttttgtgtataatcctgcataa | 50 | 95331 | 95282 | 48 | 33 | 49.5 | 15 | 157 | 118 | 0.4725 | 0.156 | 3.028846154 |
| ndhB-25 | taataacttgattatttttagataatagtagatagaaagaacgctcgtaa | 50 | 95167 | 95118 | 47 | 33 | 54 | 22 | 220 | 186 | 0.439 | 0.134 | 3.276119403 |
| ndhB-26 | ctagaagctaaaaagggtatcctgagcaatcgcaataatcgggttcattg | 50 | 94990 | 94941 | 48 | 33 | 49.5 | 15 | 171 | 116 | 0.4785 | 0.205 | 2.334146341 |
| ndhB-27 | cattttgggcggaaccgatctactaattctttgattccagttagtattag | 50 | 94910 | 94861 | 48 | 33 | 59.5 | 25 | 221.5 | 218 | 0.425 | 0.132 | 3.21969697 |
| ndhF-20 | ctcgaagtttccttcgtaacccccggaacttcttcgtagtggctccgttc | 50 | 109146 | 109195 | 48 | 30 | 143 | 67.5 | 1144.5 | 849 | 0.1665 | 0.07 | 2.378571429 |
| ndhF-19 | tgtgatggtttttcaatcttttatactaggtaatctagtatccttatgca | 50 | 109404 | 109453 | 48 | 30 | 38.5 | 16.5 | 187.5 | 136.5 | 0.3375 | 0.129 | 2.61627907 |
| ndhF-18 | accacattttccatagggccctcttatctcttccttctccgagctcgggt | 50 | 109504 | 109553 | 45 | 24 | 19 | 16 | 95 | 75 | 0.315 | 0.2145 | 1.468531469 |
| ndhF-17 | gttttattgcgggacagctcatgatgttcatatcgatctattatgcgcct | 50 | 109604 | 109653 | 48 | 30 | 29 | 13 | 173 | 133 | 0.307 | 0.1235 | 2.48582996 |
| ndhF-16 | gtatcttttgtttcatttcttctggaacaatcacaaacactttttgatt | 50 | 109704 | 109753 | 48 | 30 | 38.5 | 17 | 164 | 139 | 0.313 | 0.1365 | 2.293040293 |
| ndhF-15 | ttcctgaataatctcatttttcaattattcaaccatttcattttaccaag | 50 | 109804 | 109853 | 48 | 30 | 34.5 | 18.5 | 135 | 111 | 0.3615 | 0.178 | 2.030898876 |
| ndhF-VL | agtcaacatttatatgtttcgatgcaacaacaagatgttatttgtaacaa | 50 | 109872 | 109921 | 48 | 30 | 31.5 | 11 | 186 | 99.5 | 0.3505 | 0.178 | 1.969101124 |
| ndhF-14 | agatgttatttgtaacaagtagttttgttggttggttaattggtcacatt | 49 | 109904 | 109953 | 48 | 33 | 107.5 | 25 | 515.5 | 258 | 0.434 | 0.106 | 4.094339623 |
| ndhF-mature | gtagttttgttggttggttaattggtcacattttattcatgaaatgggtt | 50 | 109922 | 109971 | 48 | 30 | 72 | 16 | 298.5 | 150.5 | 0.341 | 0.142 | 2.401408451 |
| ndhF-13 | gcaaaataattctattaggtctaatgtagttattagatctaataagtat | 49 | 109995 | 110044 | 48 | 33 | 53 | 22 | 170 | 126 | 0.409 | 0.172 | 2.377906977 |
| ndhF-12 | ctcttatttattacctgtgtctactatttaggcagaataccatcacccat | 50 | 110095 | 110144 | 48 | 33 | 110.5 | 29 | 478.5 | 471 | 0.2815 | 0.1 | 2.815 |
| ndhF-11 | accaagaggtatccaccgaagaagctccttttccttctctttttcggaa | 50 | 110195 | 110244 | 48 | 32 | 300.5 | 151.5 | 3364.5 | 3193 | 0.1515 | 0.0555 | 2.72972973 |
| ndhF-10 | tggaaaagacaaaattaataaggatgatgaattccacgttcgaacatact | 50 | 110295 | 110344 | 48 | 33 | 67.5 | 24 | 462.5 | 344 | 0.285 | 0.077 | 3.701298701 |
| ndhF-9 | aagaaaattcgaatttagaatttttcaaaataaaaaaaaaagaggatcat | 50 | 110384 | 110433 | 36 | 24 | 26 | 9 | 63.5 | 54.5 | 0.5025 | 0.3325 | 1.511278195 |
| ndhF-8 | aattaagtaaggttaaatttaataaagatgaataaaaaggcttatataaa | 50 | 110724 | 110773 | 48 | 33 | 35 | 10 | 139.5 | 85 | 0.531 | 0.218 | 2.435779817 |

| | | | | | | | | | | | | | |
|---|---|---|---|---|---|---|---|---|---|---|---|---|---|
| ndhF-7 | tttttattaagtttttttttcttcttctttaccccataaagagagtgaata | 50 | 111170 | 111219 | 47 | 32 | 36 | 11.5 | 124 | 81.5 | 0.5 | 0.2405 | 2.079002079 |
| ndhF-6 | gatcctaaaaacaacaaagctttcgaataagcatgagtaatcaaatgaa | 49 | 111560 | 111609 | 48 | 33 | 54 | 14 | 251.5 | 127 | 0.4155 | 0.152 | 2.733552632 |
| ndhF-5 | atagaaatgcacacaaagtaaggaataagagatttactctattatttaaa | 50 | 111942 | 111991 | 48 | 33 | 44 | 10 | 207 | 94 | 0.4255 | 0.191 | 2.227748691 |
| ndhF-4 | acatataattgtcactataaatcagaaccaaaatcccaacagttgtaatt | 50 | 112293 | 112342 | 46 | 33 | 34 | 13 | 137.5 | 61 | 0.5525 | 0.294 | 1.879251701 |
| ndhF-3 | tgatccatgaatattgatatgtatgttccataaaataaaaaatccttttt | 50 | 112608 | 112657 | 43 | 33 | 26 | 11 | 95 | 77 | 0.433 | 0.286 | 1.513986014 |
| ndhF-2 | ttgtgattttatgcaaatacagaaaaagtgaattataattccatattaca | 50 | 112909 | 112958 | 48 | 30 | 18 | 7 | 85 | 66 | 0.3905 | 0.234 | 1.668803419 |
| psbD-prom | ctttttttgcgtgtattaaatacttgtctattaaatactatagtatcaact | 50 | | | 48 | 30 | 475 | 26.5 | 747.5 | 640.5 | 0.723 | 0.0565 | 12.79646018 |
| psbD-IG | tggtaaatttaccaagggctatagtcattagcgatcctcctattc | 45 | | | 48 | 30 | 694 | 52 | 1113.5 | 935.5 | 0.7625 | 0.0685 | 11.13138686 |
| psbD2 | gcaccatgaatagcgcatagcagagccgcgcccagtacaccggcgactcc | 50 | 121119 | 121168 | 48 | 30 | 305.5 | 46 | 688 | 703.5 | 0.762 | 0.073 | 10.43835616 |
| psbD3 | gtatagaaagtctcaaattctggatcttccgctgcacggatttcctggga | 50 | 120819 | 120868 | 48 | 30 | 1347 | 115 | 2532 | 2231.5 | 0.769 | 0.072 | 10.68055556 |
| psbD-psbC | agctaaagttaaagagcgtttccacgtggtagaacctcctcagggaatat | 50 | | | 48 | 30 | 749 | 83 | 1106.5 | 1038.5 | 0.7875 | 0.081 | 9.722222222 |
| psaJ | aatgcatctggaaataaacgattaatctctattaataaacctgctaacga | 50 | 87441 | 87490 | 48 | 33 | 338.5 | 90 | 4413.5 | 2386 | 0.134 | 0.072 | 1.861111111 |
| trnF | atttgaactggtgacacgaggattttcagtcctctgctctaccaactgag | 50 | 106245 | 106294 | 48 | 33 | 245 | 868 | 15360 | 18463 | 0.041 | 0.066 | 0.621212121 |
| trnT | | | | | 48 | 30 | 234 | 294.5 | 6785 | 5507 | 0.0545 | 0.068 | 0.801470588 |
| trnL(CAA)_s | | | | | 48 | 30 | 537 | 991 | 20361.5 | 22864 | 0.039 | 0.044 | 0.886363636 |

[1] oligo nucleotide genome position in NC_000932.1

[2] each probe is spotted with 12 replications per chip; spots with less than 5 detectable spots were removed from the data set

[3] the median of red (635 nm) or green (532 nm) fluorescence, from the replicates of each probe is calculated, including a local background substraction, performed by the GenePix Pro 6.0

[4] median of ratios of the background red (635 nm) and green (532 nm) fluorescence

[5] differential enrichment of IP over control- IP, based on the median of ratios of red (635 nm) or green (532 nm) fluorescence
