## Supplemental Table 3 for "The chloroplast ribonucleoprotein CP33B quantitatively binds the *psbA* mRNA"

**Supplemental Table S3:** summary of sites in the chloroplast genome of Arabidopsis that correspond to the top motif of the RBNS analysis.

| | target sites | | |
|---|---|---|---|
| RBNS motif [1] | *psbA* CDS[2] | *psbA;* non-translated[2;3] | other [No] |
| GGUUACU | / | / | 16 |
| GGCUAUU | / | / | 24 |
| GGCUACU | / | / | 10 |
| GGUUAUU | 688-682 | / | 40 |

[1] The RBNS top motif was further separated into four distinct sequences based on C/U ambiguities at position 3 and 6.

[2] target site position within the Arabidopsis thaliana chloroplast genome (NC_000932.1)

[3] target sites in the intergenic regions between *trnK* and *psbA* and between *trnH* and *psbA.*
