## Supplemental Table 4 for "The chloroplast ribonucleoprotein CP33B quantitatively binds the *psbA* mRNA"

**Supplemental Table 4**: RIP-chip data for Fig. 5B (CP33B RIP-chip from membranes)

| Name | | | Median(F532 Median - B532) | | Median (Median of Ratios (635/532)) | | Diff. Enrich. [2] |
|---|---|---|---|---|---|---|---|
| | Spot Count CP33B [1] | Spot Count control [1] | CP33B | control | CP33B | control | (CP33B/ control) |
| psbA | 12 | 12 | 4457.5 | 5758 | 2.3667 | 0.068 | 34.80441176 |
| psbA-Kex2 | 12 | 12 | 864 | 979.5 | 0.9863 | 0.13 | 7.586923077 |
| Kex2-matK | 12 | 12 | 836.5 | 628.5 | 0.5026 | 0.277 | 1.814440433 |
| matK-1 | 12 | 12 | 1170.5 | 648.5 | 0.5852 | 0.3405 | 1.718649046 |
| matK-2 | 12 | 12 | 1643.5 | 1407.5 | 1.0892 | 0.4695 | 2.319914803 |
| Kint | 12 | 12 | 1036.5 | 670.5 | 0.8155 | 0.3675 | 2.219047619 |
| Kint-Kex1 | 12 | 12 | 597.5 | 506 | 0.6818 | 0.3815 | 1.787155963 |
| IG_K-rps16 | 12 | 12 | 1451.5 | 799.5 | 0.8372 | 0.4115 | 2.034507898 |
| rps16ex2 | 12 | 12 | 1752.5 | 1322 | 1.1431 | 0.4855 | 2.354479918 |
| rps16ex2-in-ex1 | 12 | 12 | 1044.5 | 761 | 0.6769 | 0.397 | 1.705037783 |
| rps16in-1 | 12 | 12 | 1048.5 | 697 | 0.9583 | 0.4455 | 2.151066218 |
| rps16in-2 | 12 | 12 | 712.5 | 474 | 0.8218 | 0.412 | 1.994660194 |
| rps16ex2-IG | 12 | 12 | 1419 | 1344.5 | 0.7427 | 0.4125 | 1.800484848 |
| Q | 12 | 12 | 7077 | 2449 | 0.1526 | 0.089 | 1.714606742 |
| psbI-S | 12 | 12 | 1477 | 1394 | 0.4249 | 0.1765 | 2.407365439 |
| IG-psbI-G-1 | 12 | 12 | 16009.5 | 14259 | 0.0546 | 0.031 | 1.761290323 |
| IG-psbI-G-2 | 12 | 12 | 2110 | 1676.5 | 0.3409 | 0.152 | 2.242763158 |
| Gex1-in | 12 | 12 | 5869 | 3894 | 0.3486 | 0.098 | 3.557142857 |
| in-Gex2 | 12 | 12 | 3558 | 2900 | 0.3059 | 0.117 | 2.614529915 |
| Gex2-R | 12 | 12 | 2145.5 | 1063 | 0.7014 | 0.2405 | 2.916424116 |
| atpA-1 | 12 | 12 | 1168 | 840.5 | 0.9646 | 0.4375 | 2.2048 |
| atpA-3 | 12 | 12 | 573 | 383 | 0.8008 | 0.296 | 2.705405405 |
| atpA-atpFex2 | 12 | 12 | 645.5 | 646 | 0.8015 | 0.284 | 2.822183099 |
| atpFex2-in | 11 | 12 | 1548 | 1205 | 1.0262 | 0.2915 | 3.520411664 |
| atpFex2-in-ex1 | 12 | 12 | 2410 | 2201.5 | 0.707 | 0.2725 | 2.594495413 |
| atpFin-ex1 | 12 | 12 | 989 | 782 | 0.7371 | 0.2995 | 2.461101836 |
| atpFex1-IG | 11 | 12 | 1902 | 2310.5 | 0.8652 | 0.272 | 3.180882353 |
| atpH | 12 | 12 | 882.5 | 743.5 | 0.8113 | 0.3215 | 2.52348367 |
| atpH-IG | 12 | 12 | 6638 | 8399 | 1.1746 | 0.229 | 5.129257642 |
| IG-atpI | 12 | 12 | 1084.5 | 1240.5 | 0.7973 | 0.339 | 2.351917404 |
| atpI | 12 | 12 | 1248 | 954 | 0.8526 | 0.3045 | 2.8 |
| atpI-rps2 | 12 | 12 | 2089 | 2330 | 0.8645 | 0.3 | 2.881666667 |
| rps2 | 12 | 12 | 1651 | 1088.5 | 0.7665 | 0.38 | 2.017105263 |
| rps2-IG | 11 | 12 | 859 | 768.5 | 0.5698 | 0.377 | 1.511405836 |
| rpoC2-1 | 12 | 12 | 1239.5 | 1036 | 0.6958 | 0.36 | 1.932777778 |
| rpoC2-2 | 10 | 12 | 1298 | 810 | 0.5502 | 0.352 | 1.563068182 |
| rpoC2-3 | 12 | 9 | 1650.5 | 274 | 0.5761 | 0.396 | 1.45479798 |
| rpoC2-4 | 12 | 12 | 1585 | 980 | 0.5131 | 0.3695 | 1.388633288 |
| rpoC2-5 | 12 | 12 | 1038 | 804 | 0.56 | 0.3745 | 1.495327103 |
| rpoC2-6 | 12 | 12 | 1752.5 | 1123 | 0.511 | 0.3445 | 1.483309144 |

| | | | | | | | |
|---|---|---|---|---|---|---|---|
| rpoC2-rpoC1-2 | 12 | 12 | 820 | 621.5 | 0.5082 | 0.3835 | 1.325162973 |
| rpoC1ex1-1 | 12 | 12 | 1645 | 1042.5 | 0.5278 | 0.362 | 1.45801105 |
| rpoC1ex1-2 | 12 | 12 | 1093.5 | 833.5 | 0.4179 | 0.3325 | 1.256842105 |
| rpoC1ex1-in | 12 | 12 | 873 | 564.5 | 0.5803 | 0.3625 | 1.600827586 |
| rpoC1ex1-in-ex2 | 12 | 12 | 1454 | 937.5 | 0.5397 | 0.369 | 1.462601626 |
| rpoC1in | 12 | 12 | 1294.5 | 885.5 | 0.518 | 0.3615 | 1.432918396 |
| rpoC1in-ex2 | 12 | 12 | 1348.5 | 991.5 | 0.6062 | 0.4045 | 1.498640297 |
| rpoC1-rpoB-1 | 12 | 12 | 1960.5 | 1337 | 0.5579 | 0.4245 | 1.314252061 |
| rpoC1-rpoB-2 | 12 | 12 | 2637.5 | 1645 | 0.5992 | 0.371 | 1.61509434 |
| rpoB-1 | 12 | 12 | 1306 | 887 | 0.3346 | 0.315 | 1.062222222 |
| rpoB-2 | 12 | 12 | 454 | 467.5 | 0.5264 | 0.3355 | 1.56900149 |
| rpoB-3 | 12 | 12 | 1925 | 1180 | 0.8379 | 0.4415 | 1.897848245 |
| rpoB-4 | 12 | 12 | 1035 | 577.5 | 0.5558 | 0.429 | 1.295571096 |
| rpoB-IG | 12 | 12 | 1195 | 952.5 | 0.6741 | 0.4415 | 1.526840317 |
| IG rpoB-C | 12 | 12 | 1253.5 | 1271 | 0.6566 | 0.4295 | 1.528754366 |
| IG-C | 12 | 12 | 4520 | 3696 | 0.1813 | 0.095 | 1.908421053 |
| C-IG | 12 | 12 | 374 | 383 | 0.4508 | 0.3775 | 1.194172185 |
| IG-petN | 12 | 12 | 1517.5 | 1083 | 0.4802 | 0.257 | 1.86848249 |
| petN-IG | 12 | 12 | 255.5 | 279.5 | 0.5782 | 0.325 | 1.779076923 |
| psbM | 12 | 12 | 477.5 | 309 | 0.63 | 0.3565 | 1.767180926 |
| IG psbM-D-1 | 12 | 12 | 829.5 | 574.5 | 0.5537 | 0.339 | 1.633333333 |
| IG psbM-D-2 | 12 | 12 | 1017.5 | 541.5 | 0.7224 | 0.4405 | 1.639954597 |
| D | 12 | 12 | 1094.5 | 1262.5 | 0.2814 | 0.1625 | 1.731692308 |
| Y | 12 | 12 | 644 | 436 | 0.6489 | 0.3145 | 2.06327504 |
| E | 12 | 12 | 1839 | 1423 | 0.7644 | 0.344 | 2.222093023 |
| IG E-T | 12 | 12 | 1399.5 | 923.5 | 0.8162 | 0.4505 | 1.811764706 |
| T | 12 | 12 | 941.5 | 982.5 | 0.5201 | 0.246 | 2.114227642 |
| IG T-psbD-1 | 12 | 11 | 1835.5 | 1808 | 0.6405 | 0.289 | 2.216262976 |
| IG T-psbD-2 | 12 | 12 | 2834 | 1580.5 | 0.9905 | 0.264 | 3.751893939 |
| psbD-1 | 12 | 12 | 3610.5 | 2070.5 | 1.6485 | 0.354 | 4.656779661 |
| psbD-2 | 12 | 11 | 1242 | 1198 | 0.6895 | 0.228 | 3.024122807 |
| psbD-psbC | 12 | 12 | 767.5 | 807 | 0.7882 | 0.2515 | 3.133996024 |
| psbC-1 | 12 | 12 | 1182.5 | 1291.5 | 0.4648 | 0.168 | 2.766666667 |
| psbC-2 | 12 | 12 | 2474.5 | 2403.5 | 0.7833 | 0.171 | 4.580701754 |
| psbC-S | 12 | 12 | 2126 | 2261.5 | 0.6433 | 0.2055 | 3.130413625 |
| S-psbZ | 12 | 12 | 2275 | 2184 | 0.9324 | 0.311 | 2.99807074 |
| IG psbZ-G | 12 | 12 | 1567.5 | 1228.5 | 0.9597 | 0.343 | 2.797959184 |
| G-fM | 12 | 12 | 1072.5 | 933.5 | 0.2912 | 0.1405 | 2.072597865 |
| fm-rps14 | 12 | 12 | 2645.5 | 2093 | 0.182 | 0.096 | 1.895833333 |
| rps14-psaB-1 | 12 | 12 | 3391 | 2760.5 | 0.5684 | 0.3125 | 1.81888 |
| rps14-psaB-2 | 12 | 12 | 2662.5 | 2047 | 0.6741 | 0.334 | 2.018263473 |
| psaB-1 | 12 | 12 | 5786 | 5122 | 1.3097 | 0.403 | 3.249875931 |
| psaB-2 | 12 | 12 | 1296.5 | 646 | 1.1025 | 0.415 | 2.656626506 |
| psaB-psaA-1 | 12 | 11 | 1230 | 1080 | 0.9373 | 0.372 | 2.519623656 |
| psaB-psaA-2 | 12 | 12 | 1140 | 1095 | 0.8995 | 0.394 | 2.282994924 |
| psaA-1 | 12 | 12 | 4568 | 3143 | 0.8127 | 0.3335 | 2.436881559 |
| psaA-2 | 12 | 12 | 1422.5 | 1184 | 1.2012 | 0.4545 | 2.64290429 |
| psaA-3 | 12 | 12 | 4696.5 | 3557 | 1.8214 | 0.439 | 4.148974943 |

| | | | | | | | |
|---|---|---|---|---|---|---|---|
| psaA-IG-1 | 12 | 12 | 4148 | 2448 | 1.1802 | 0.363 | 3.251239669 |
| psaA-IG-2 | 12 | 12 | 863 | 601.5 | 1.1382 | 0.4465 | 2.549160134 |
| IG-ycf3ex3 | 12 | 12 | 1440 | 1223.5 | 0.9758 | 0.437 | 2.232951945 |
| ycf3ex3-in2 | 12 | 12 | 1342.5 | 1030.5 | 0.7966 | 0.4235 | 1.880991736 |
| ycf3ex3-in2-ex2 | 11 | 11 | 1004 | 720 | 0.6188 | 0.396 | 1.562626263 |
| ycf3in2 | 11 | 12 | 915 | 605 | 0.8386 | 0.424 | 1.977830189 |
| ycf3ex2 | 12 | 12 | 638.5 | 470 | 0.6433 | 0.4175 | 1.540838323 |
| ycf3in1 | 12 | 11 | 2184.5 | 1132 | 0.9436 | 0.438 | 2.1543379 |
| acf3ex1 | 12 | 12 | 1717 | 1111 | 0.7903 | 0.435 | 1.816781609 |
| S-GGA | 12 | 12 | 4125 | 4148 | 0.1365 | 0.0765 | 1.784313725 |
| rps4-1 | 12 | 12 | 1270 | 968 | 0.8183 | 0.4555 | 1.796487377 |
| rps4-2 | 12 | 12 | 866.5 | 648.5 | 0.5187 | 0.4085 | 1.269767442 |
| T-UGU | 12 | 12 | 1016.5 | 510 | 0.5201 | 0.3385 | 1.53648449 |
| T-IG | 12 | 12 | 3035.5 | 2104 | 0.1547 | 0.088 | 1.757954545 |
| L-UAA ex1-in | 12 | 12 | 5904 | 4140.5 | 0.1029 | 0.074 | 1.390540541 |
| L-UAA in-ex2 | 12 | 12 | 2197 | 2892 | 0.175 | 0.1005 | 1.741293532 |
| L-UAA ex2-IG | 12 | 12 | 4135 | 3661 | 0.3661 | 0.1985 | 1.844332494 |
| F-GAA-1 | 12 | 12 | 3121 | 2173.5 | 0.07 | 0.065 | 1.076923077 |
| F-GAA-2 | 12 | 12 | 1695 | 913.5 | 0.5481 | 0.285 | 1.923157895 |
| ndhJ | 12 | 12 | 1105.5 | 1152 | 0.6209 | 0.3225 | 1.925271318 |
| ndhJ-ndhK | 12 | 12 | 1400.5 | 1040.5 | 0.595 | 0.4015 | 1.481942715 |
| ndhK | 12 | 12 | 699 | 557 | 0.7931 | 0.4025 | 1.970434783 |
| ndhK-ndhC | 12 | 12 | 1356 | 873.5 | 0.8785 | 0.376 | 2.33643617 |
| ndhC-IG | 12 | 12 | 3661.5 | 3052.5 | 1.1648 | 0.395 | 2.948860759 |
| IG ndhC-V-UAA | 12 | 12 | 1643 | 1249.5 | 0.8967 | 0.415 | 2.160722892 |
| Vex2-in | 12 | 12 | 1222 | 824 | 0.7315 | 0.396 | 1.847222222 |
| Vin-ex1 | 12 | 12 | 2030.5 | 1418 | 0.3297 | 0.234 | 1.408974359 |
| Vex1-atpE | 12 | 12 | 3249 | 3188 | 0.1946 | 0.1035 | 1.880193237 |
| atpE-atpB | 12 | 12 | 1130 | 1177.5 | 0.4655 | 0.3085 | 1.5089141 |
| atpB-1 | 12 | 12 | 2606 | 3574.5 | 0.5138 | 0.306 | 1.679084967 |
| atpB-2 | 12 | 12 | 1508 | 1120 | 0.7651 | 0.3345 | 2.287294469 |
| atpB-3 | 12 | 12 | 2131 | 1702 | 0.581 | 0.303 | 1.917491749 |
| atpB-rbcL | 12 | 12 | 2616.5 | 2865 | 0.406 | 0.2395 | 1.69519833 |
| rbcL-1 | 12 | 12 | 9203.5 | 6701 | 0.2282 | 0.0845 | 2.700591716 |
| rbcL-2 | 12 | 12 | 1573.5 | 1632 | 0.1148 | 0.0665 | 1.726315789 |
| rbcL-3 | 12 | 12 | 7827 | 7955.5 | 0.1883 | 0.0635 | 2.965354331 |
| IG rbcL-accD | 12 | 12 | 1902.5 | 1802 | 0.5362 | 0.2945 | 1.820713073 |
| accD-1 | 12 | 12 | 1004 | 668 | 0.5684 | 0.2965 | 1.91703204 |
| accD-2 | 12 | 12 | 1818 | 1436 | 0.4893 | 0.2795 | 1.750626118 |
| accD-3 | 12 | 12 | 1365.5 | 1240.5 | 0.7315 | 0.3735 | 1.958500669 |
| IG accD-psaI | 12 | 12 | 2184 | 1587 | 0.7203 | 0.3925 | 1.835159236 |
| psaI | 12 | 12 | 1399.5 | 845 | 0.5285 | 0.327 | 1.616207951 |
| ycf4-1 | 12 | 12 | 2885.5 | 2116.5 | 0.5334 | 0.274 | 1.946715328 |
| ycf4-2 | 12 | 12 | 1570 | 1324 | 0.6223 | 0.318 | 1.956918239 |
| cemA-1 | 11 | 12 | 1437 | 1559 | 1.085 | 0.418 | 2.59569378 |
| cemA-2 | 12 | 12 | 2138.5 | 1641 | 0.9191 | 0.3605 | 2.549514563 |
| cemA-3 | 12 | 12 | 2536 | 1587.5 | 0.7322 | 0.3145 | 2.328139905 |
| petA-1 | 12 | 12 | 1126 | 905 | 0.5775 | 0.3175 | 1.818897638 |

| | | | | | | | |
|---|---|---|---|---|---|---|---|
| petA-2 | 12 | 12 | 2548.5 | 2152.5 | 0.5432 | 0.2545 | 2.134381139 |
| petA-3 | 12 | 12 | 1683.5 | 1161.5 | 0.875 | 0.4045 | 2.1631644 |
| IG petA-psbJ | 12 | 12 | 1937.5 | 1518 | 0.5705 | 0.2725 | 2.093577982 |
| psbJLF | 12 | 12 | 6727.5 | 3590 | 0.1281 | 0.1045 | 1.225837321 |
| psbFE | 12 | 12 | 3162 | 2952 | 0.3794 | 0.1695 | 2.238348083 |
| IG psbE-petL-1 | 12 | 12 | 3746.5 | 1869 | 0.5061 | 0.209 | 2.4215311 |
| IG psbE-petL-2 | 12 | 12 | 1262 | 772.5 | 0.7182 | 0.357 | 2.011764706 |
| IG psbE-petL-3 | 12 | 12 | 1510.5 | 627.5 | 0.7903 | 0.3615 | 2.186168741 |
| petL | 12 | 12 | 3732.5 | 2709 | 0.4739 | 0.2335 | 2.029550321 |
| petG-W-CCA | 12 | 12 | 2216.5 | 1493 | 0.5698 | 0.258 | 2.208527132 |
| P-UGG | 12 | 12 | 2695.5 | 2575 | 0.2765 | 0.106 | 2.608490566 |
| psaJ | 12 | 12 | 5939.5 | 4665 | 0.175 | 0.083 | 2.108433735 |
| IG psaJ-rpl33 | 12 | 12 | 3247 | 3049 | 0.7602 | 0.279 | 2.724731183 |
| rpl33 | 12 | 12 | 2269.5 | 1008.5 | 1.0598 | 0.325 | 3.260923077 |
| rpl33-rps18 | 12 | 12 | 4177.5 | 3253 | 0.9191 | 0.345 | 2.664057971 |
| rps18 | 12 | 12 | 1816.5 | 1466.5 | 0.8316 | 0.375 | 2.2176 |
| rpl20 | 12 | 12 | 1455 | 975 | 0.63 | 0.405 | 1.555555556 |
| rpl20-IG | 12 | 12 | 1363.5 | 776.5 | 0.6783 | 0.4555 | 1.489132821 |
| in1-rps12ex1 | 12 | 12 | 1374.5 | 1564 | 0.5978 | 0.364 | 1.642307692 |
| rps12ex1-clpPex3 | 12 | 12 | 2239 | 1635.5 | 0.665 | 0.299 | 2.224080268 |
| clpPex3-in2 | 12 | 12 | 2380 | 1408.5 | 0.6993 | 0.3825 | 1.828235294 |
| in2-clpPex2 | 12 | 12 | 1650.5 | 1472.5 | 0.5992 | 0.388 | 1.544329897 |
| clpPex2-in1 | 12 | 12 | 1859 | 770.5 | 0.6713 | 0.365 | 1.839178082 |
| clpPin1 | 12 | 12 | 2228.5 | 1900.5 | 0.5663 | 0.3075 | 1.841626016 |
| clpPex1 | 12 | 12 | 4715.5 | 3739.5 | 0.7588 | 0.2365 | 3.20845666 |
| psbB-1 | 12 | 12 | 7451 | 5125.5 | 0.4557 | 0.168 | 2.7125 |
| psbB-2 | 12 | 12 | 1763.5 | 1683 | 0.3297 | 0.169 | 1.950887574 |
| psbBTN | 11 | 12 | 1216 | 1839 | 0.686 | 0.2385 | 2.876310273 |
| psbNH | 12 | 12 | 2911 | 2097 | 0.2695 | 0.114 | 2.364035088 |
| psbH-petBex1-in | 12 | 12 | 2030.5 | 1380 | 0.5859 | 0.2985 | 1.96281407 |
| petBin | 12 | 12 | 2083.5 | 1582 | 1.1102 | 0.433 | 2.563972286 |
| petBin-ex2 | 12 | 12 | 5248.5 | 3805 | 0.8302 | 0.195 | 4.257435897 |
| petBex2 | 12 | 12 | 2270 | 2036.5 | 0.3472 | 0.1615 | 2.149845201 |
| petBex2-petDex1-in | 12 | 12 | 2784.5 | 2435.5 | 0.5187 | 0.2015 | 2.574193548 |
| petDin | 12 | 12 | 2353.5 | 1589 | 0.6314 | 0.279 | 2.263082437 |
| petDex2 | 12 | 12 | 2714 | 2913.5 | 0.7511 | 0.189 | 3.974074074 |
| petD-rpoA | 12 | 12 | 1002 | 915 | 0.5544 | 0.316 | 1.75443038 |
| rpoA | 11 | 12 | 2050 | 1491.5 | 0.693 | 0.3655 | 1.896032832 |
| rpoA-rps11 | 12 | 12 | 1770 | 1037.5 | 0.6503 | 0.3785 | 1.718097754 |
| rps11-rpl36 | 12 | 12 | 1898.5 | 794.5 | 0.6657 | 0.3735 | 1.782329317 |
| IG rpl36-rps8 | 12 | 12 | 977 | 1226 | 0.7455 | 0.3725 | 2.001342282 |
| rps8 | 12 | 12 | 1092 | 1409 | 0.7987 | 0.4425 | 1.804971751 |
| rps8-rpl14 | 12 | 12 | 2448.5 | 1729 | 0.8554 | 0.386 | 2.216062176 |
| rpl14 | 12 | 12 | 1683.5 | 1318 | 0.7007 | 0.336 | 2.085416667 |
| rpl16ex2 | 12 | 12 | 1151.5 | 790.5 | 0.7007 | 0.4005 | 1.749563046 |
| rpl16in-1 | 12 | 12 | 3242 | 2742.5 | 0.9037 | 0.4155 | 2.174969916 |

| | | | | | | | |
|---|---|---|---|---|---|---|---|
| rpl16in-2 | 12 | 12 | 825 | 440 | 0.8316 | 0.423 | 1.965957447 |
| rpl16in-ex1 | 12 | 12 | 1895 | 1703 | 0.8323 | 0.4475 | 1.859888268 |
| rps3-1 | 12 | 12 | 2257 | 1396 | 0.7546 | 0.4225 | 1.786035503 |
| rps3-2 | 12 | 12 | 1431.5 | 1398.5 | 0.7518 | 0.432 | 1.740277778 |
| rpl22 | 11 | 12 | 1656 | 1340.5 | 0.945 | 0.4425 | 2.13559322 |
| rps19 | 12 | 12 | 3028.5 | 1539 | 0.9317 | 0.426 | 2.187089202 |
| rps19-rpl2ex2 | 12 | 12 | 2545.5 | 2155 | 0.8204 | 0.3595 | 2.282058414 |
| rpl2ex2-in | 12 | 12 | 2578 | 2114 | 0.7406 | 0.363 | 2.040220386 |
| rpl2ex2-in-ex1 | 12 | 12 | 2854 | 2053 | 0.7077 | 0.369 | 1.917886179 |
| rpl2in-ex1 | 12 | 12 | 2105 | 1533 | 0.7028 | 0.362 | 1.941436464 |
| rpl2ex1 | 12 | 12 | 2838 | 2029.5 | 0.7238 | 0.329 | 2.2 |
| rpl23-I-CAU | 12 | 12 | 2486.5 | 1689.5 | 0.8267 | 0.41 | 2.016341463 |
| I-CAU-ycf2 | 12 | 12 | 2637 | 1642.5 | 0.4186 | 0.251 | 1.667729084 |
| ycf2-1 | 12 | 12 | 2424.5 | 1668 | 0.4998 | 0.344 | 1.452906977 |
| ycf2-3 | 12 | 12 | 2527 | 1768.5 | 0.5677 | 0.3765 | 1.507835325 |
| ycf2-4 | 12 | 12 | 2806.5 | 1767 | 0.5663 | 0.3625 | 1.562206897 |
| ycf2-5 | 12 | 12 | 1930 | 1134.5 | 0.4816 | 0.3905 | 1.233290653 |
| ycf2-6 | 12 | 12 | 2054.5 | 1384 | 0.4004 | 0.3195 | 1.253208138 |
| ycf2-7 | 12 | 12 | 2576 | 1797 | 0.4298 | 0.348 | 1.235057471 |
| ycf2-8 | 12 | 12 | 2068.5 | 1238 | 0.6727 | 0.385 | 1.747272727 |
| ycf2-IG | 12 | 12 | 1945 | 1308.5 | 0.5103 | 0.3805 | 1.341130092 |
| IG-L-CAA | 12 | 12 | 3199 | 1409 | 0.532 | 0.4265 | 1.247362251 |
| L-CAA | 12 | 12 | 1960 | 1465.5 | 0.4032 | 0.3535 | 1.140594059 |
| ndhBex2-1 | 12 | 12 | 2714 | 1510.5 | 0.5558 | 0.3375 | 1.646814815 |
| ndhBex2-2 | 12 | 12 | 1494 | 912.5 | 0.4935 | 0.3175 | 1.554330709 |
| ndhBex2-in | 12 | 12 | 1860.5 | 1010 | 0.6237 | 0.384 | 1.62421875 |
| ndhBex2-in-ex1 | 12 | 12 | 3609 | 2210.5 | 0.595 | 0.392 | 1.517857143 |
| ndhBin-ex1 | 12 | 12 | 1115.5 | 818.5 | 0.5866 | 0.3515 | 1.668847795 |
| ndhBex1-1 | 12 | 12 | 2626 | 1828.5 | 0.63 | 0.3615 | 1.742738589 |
| ndhBex1-2 | 12 | 12 | 2916.5 | 1806.5 | 0.6923 | 0.3245 | 2.133436055 |
| IG-rps7 | 12 | 12 | 1529 | 1031.5 | 0.525 | 0.3585 | 1.464435146 |
| rps7-rps12ex3 | 12 | 12 | 2738.5 | 2349.5 | 0.4809 | 0.293 | 1.641296928 |
| rps12in2-ex2 | 12 | 12 | 2918 | 1351.5 | 0.4886 | 0.338 | 1.44556213 |
| rps12ex2 | 12 | 12 | 2611.5 | 1379 | 0.4627 | 0.301 | 1.537209302 |
| rps12in1 | 12 | 12 | 3528 | 2074.5 | 0.4893 | 0.3275 | 1.494045802 |
| IG rps12-V-GAC-1 | 12 | 12 | 3616.5 | 1157.5 | 0.4872 | 0.288 | 1.691666667 |
| IG rps12-V-GAC-2 | 12 | 12 | 1473.5 | 856.5 | 0.4081 | 0.305 | 1.338032787 |
| IG rps12-V-GAC-3 | 11 | 12 | 1989 | 1373 | 0.3892 | 0.2655 | 1.465913371 |
| IG rps12-V-GAC-4 | 11 | 11 | 1855 | 1190 | 0.3808 | 0.306 | 1.244444444 |
| IG rps12-V-GAC-5 | 12 | 12 | 1903 | 1078.5 | 0.3276 | 0.252 | 1.3 |
| V-GAC | 12 | 12 | 4210 | 1512.5 | 0.3486 | 0.2175 | 1.602758621 |
| V-GAC-16S | 12 | 12 | 20754 | 15002 | 0.0245 | 0.0355 | 0.690140845 |
| 16S-1 | 12 | 12 | 37694 | 19897 | 0.035 | 0.0495 | 0.707070707 |
| 16S-2 | 12 | 12 | 13777.5 | 7170.5 | 0.0336 | 0.057 | 0.589473684 |
| 16S-3 | 12 | 12 | 12399.5 | 8502.5 | 0.0476 | 0.066 | 0.721212121 |

| | | | | | | | |
|---|---|---|---|---|---|---|---|
| 16S-4 | 12 | 12 | 11934 | 7914 | 0.0385 | 0.0555 | 0.693693694 |
| 16S-5 | 12 | 12 | 3765 | 2901.5 | 0.0987 | 0.0565 | 1.746902655 |
| I-GAU ex1 | 12 | 12 | 4515.5 | 2747 | 0.1505 | 0.0795 | 1.893081761 |
| I-GAU ex1-in | 12 | 12 | 1199.5 | 759 | 0.4165 | 0.1675 | 2.486567164 |
| I-GAU ex2 | 10 | 12 | 4034.5 | 2970.5 | 0.1708 | 0.0825 | 2.07030303 |
| I-GAU-A-UGC | 12 | 12 | 1172.5 | 793 | 0.1848 | 0.1305 | 1.416091954 |
| A-UGC ex1-in | 12 | 12 | 1342.5 | 589 | 0.3234 | 0.184 | 1.757608696 |
| A-UGC -in | 12 | 12 | 2122 | 1153 | 0.2779 | 0.1705 | 1.629912023 |
| A-UGC ex1 | 12 | 12 | 2048.5 | 843 | 0.21 | 0.185 | 1.135135135 |
| A-UGC-23S | 12 | 12 | 15389.5 | 11385.5 | 0.0245 | 0.0395 | 0.620253165 |
| 23S-1 | 12 | 12 | 11735.5 | 7756.5 | 0.0203 | 0.0315 | 0.644444444 |
| 23S-2 | 12 | 12 | 3026.5 | 2855 | 0.0287 | 0.048 | 0.597916667 |
| 23S-3 | 12 | 12 | 15407 | 11179 | 0.0252 | 0.04 | 0.63 |
| 23S-4 | 12 | 12 | 22195.5 | 18027 | 0.0287 | 0.0455 | 0.630769231 |
| 23S-5 | 12 | 12 | 12708 | 10437 | 0.0252 | 0.0325 | 0.775384615 |
| 23S-6 | 12 | 12 | 4119 | 2794 | 0.0476 | 0.055 | 0.865454545 |
| 23S-7 | 12 | 12 | 17286.5 | 15468 | 0.0329 | 0.0555 | 0.592792793 |
| 23S-8 | 12 | 12 | 6883.5 | 6379.5 | 0.0301 | 0.0525 | 0.573333333 |
| 23S-9 | 12 | 12 | 8054 | 7717 | 0.0252 | 0.0265 | 0.950943396 |
| 4,5S | 12 | 12 | 6573 | 5012.5 | 0.0553 | 0.044 | 1.256818182 |
| R-ACG-1 | 12 | 12 | 3773.5 | 2478 | 0.161 | 0.1335 | 1.205992509 |
| R-ACG-2 | 12 | 12 | 1954 | 1027 | 0.5187 | 0.4145 | 1.251387214 |
| N-GUU-1 | 12 | 12 | 701.5 | 553.5 | 0.4032 | 0.277 | 1.455595668 |
| IG-ycf1 | 12 | 12 | 2871 | 1672.5 | 0.4326 | 0.267 | 1.620224719 |
| 3'ndhF(ycf1) | 12 | 12 | 2273.5 | 1209.5 | 0.8162 | 0.476 | 1.714705882 |
| ndhF-1 | 12 | 12 | 2472.5 | 1336.5 | 0.9947 | 0.466 | 2.134549356 |
| ndhF-2 | 12 | 12 | 2763 | 1915.5 | 0.7959 | 0.458 | 1.737772926 |
| ndhF-3 | 12 | 12 | 1828 | 1446 | 0.7749 | 0.3935 | 1.969250318 |
| ndhF-4 | 12 | 12 | 2092.5 | 1689 | 0.9625 | 0.514 | 1.872568093 |
| ndhF-5 | 12 | 12 | 2118 | 1142 | 0.7518 | 0.4025 | 1.867826087 |
| ndhF-IG | 12 | 12 | 385 | 445 | 1.0346 | 0.4165 | 2.484033613 |
| rpl32 | 12 | 12 | 1121.5 | 429.5 | 0.4683 | 0.188 | 2.490957447 |
| IG rpl32-ccsA | 12 | 12 | 740.5 | 358.5 | 0.8421 | 0.5085 | 1.656047198 |
| IG-ccsA | 12 | 12 | 596 | 363.5 | 0.6069 | 0.414 | 1.465942029 |
| ccsA | 12 | 12 | 2439 | 3015 | 0.819 | 0.3615 | 2.265560166 |
| ccsA-ndhD | 12 | 12 | 1055.5 | 698 | 0.8722 | 0.527 | 1.655028463 |
| ndhD-1 | 12 | 12 | 1038.5 | 789 | 0.7238 | 0.4365 | 1.658190149 |
| ndhD-2 | 12 | 12 | 764 | 440.5 | 0.819 | 0.4585 | 1.786259542 |
| ndhD-psaC | 12 | 12 | 1946 | 1488.5 | 1.0458 | 0.4395 | 2.379522184 |
| psaC-ndhE | 12 | 12 | 4656.5 | 3648 | 0.6139 | 0.257 | 2.388715953 |
| ndhE | 12 | 12 | 1292 | 728 | 0.847 | 0.454 | 1.865638767 |
| IG-ndhG | 12 | 12 | 2647.5 | 1725.5 | 0.7917 | 0.401 | 1.974314214 |
| ndhG | 12 | 12 | 1738 | 1141 | 0.6909 | 0.3065 | 2.254159869 |
| ndhG-ndhI | 12 | 12 | 2843.5 | 2681 | 0.9065 | 0.418 | 2.168660287 |
| ndhI | 12 | 12 | 1164.5 | 773.5 | 0.7819 | 0.418 | 1.870574163 |
| ndhI-ndhAex2 | 12 | 12 | 826 | 863 | 1.0003 | 0.4835 | 2.068872802 |
| ndhAex2-in | 12 | 12 | 862 | 685 | 0.7217 | 0.4715 | 1.530646872 |
| ndhAex2-in-ex1 | 12 | 12 | 1519.5 | 1302 | 0.5964 | 0.3975 | 1.500377358 |

| | | | | | | | |
|---|---|---|---|---|---|---|---|
| ndhAin-1 | 12 | 12 | 1394.5 | 878 | 0.8862 | 0.5585 | 1.586750224 |
| ndhAin-2 | 12 | 12 | 786.5 | 467 | 0.7784 | 0.5335 | 1.459044049 |
| ndhAin-ex1 | 12 | 12 | 962 | 690 | 0.7119 | 0.4805 | 1.481581686 |
| ndhAex1-ndhH | 12 | 12 | 1754.5 | 1411 | 1.064 | 0.515 | 2.066019417 |
| ndhH-1 | 12 | 12 | 957.5 | 665.5 | 0.7063 | 0.4595 | 1.53710555 |
| ndhH-2 | 12 | 12 | 617 | 402.5 | 0.9653 | 0.5915 | 1.631952663 |
| rps15 | 12 | 12 | 1145.5 | 774 | 0.9037 | 0.5265 | 1.71642925 |
| rps15-IG | 12 | 12 | 902 | 667.5 | 0.9212 | 0.5125 | 1.797463415 |
| ycf1-1 | 12 | 12 | 1188.5 | 723 | 0.9583 | 0.516 | 1.857170543 |
| ycf1-2 | 12 | 12 | 1332.5 | 837 | 0.9079 | 0.5275 | 1.721137441 |
| ycf1-3 | 12 | 12 | 2080.5 | 1248.5 | 1.2474 | 0.645 | 1.933953488 |
| ycf1-4 | 12 | 12 | 1535 | 954 | 0.9744 | 0.559 | 1.743112701 |
| ycf1-5 | 12 | 12 | 1545.5 | 791 | 1.0619 | 0.632 | 1.680221519 |
| ycf1-6 | 12 | 12 | 1714.5 | 1005.5 | 0.9975 | 0.575 | 1.734782609 |
| ycf1-7 | 12 | 12 | 1457 | 1026.5 | 0.7182 | 0.4825 | 1.488497409 |
| ycf1-8 | 12 | 12 | 1543 | 1147.5 | 0.6475 | 0.446 | 1.451793722 |
| ycf1-9 | 12 | 12 | 2504.5 | 1031 | 0.6524 | 0.4605 | 1.416720955 |
| N-GUU-2 | 12 | 12 | 3482 | 2457 | 0.2989 | 0.1755 | 1.703133903 |
| IG N-R | 12 | 12 | 1909 | 1423 | 0.6146 | 0.4665 | 1.317470525 |
| ycf2-9 | 12 | 12 | 1985.5 | 1307 | 0.5684 | 0.3715 | 1.530013459 |
| ycf2-I-CAU | 12 | 12 | 3249 | 1369 | 0.3997 | 0.265 | 1.508301887 |
| 18S | 12 | 12 | 2193 | 2103 | 0.0574 | 0.061 | 0.940983607 |

[1] each probe is spotted with 12 replications per chip; spots with less than 5 detectable spots were removed from the data

[2] differential enrichment of IP over control- IP, based on the median of ratios of red (635 nm) or green (532 nm) fluoresc

a set
ence
